# Dynamic Transcriptomic Remodeling in Human Neural Progenitor Cells Reveals Mechanisms for Vision Preservation in Retinitis Pigmentosa Model

**DOI:** 10.1101/2025.01.06.631620

**Authors:** Saba Shahin, Shaughn Bell, Bin Lu, Somanshu Banerjee, Vivek Swarup, Hui Xu, Jason Chetsawang, Stephany Ramirez, Jorge S Alfaro, Alexander Laperle, Soshana Svendsen, Clive N. Svendsen, Shaomei Wang

## Abstract

Human neural progenitor cells (hNPCs) have shown promise in slowing down retinal degeneration in animal models and are currently being tested in clinical trials for treating retinitis pigmentosa (RP). However, the status of grafted hNPCs and their interaction with host retinal cells over time is largely unknown. Here, we investigated single-cell transcriptomic changes in grafted hNPCs and host retinal cells following injection into a rodent model for RP. Grafted hNPCs and host retinal cells undergo dynamic transcriptomic changes in the degenerative retinal environment. Grafted hNPCs protect vision through multiple mechanisms, including trophic factor support, modulation of metabolic activity, reduction of apoptosis, oxidative stress, and inflammation, alongside extracellular matrix remodeling. CellChat analysis revealed a progressive decline in intercellular signaling and communication strength between hNPCs and host retinal cells over time. This study indicates that enhancing trophic factor supports and improving host retinal environment are key targets to enable long-term vision preservation.

## INTRODUCTION

Retinitis pigmentosa (RP) is a group of inherited retinal diseases characterized by progressive vision loss due to mutations in photoreceptors and retinal pigment epithelium (RPE). More than 80 genes with over 3000 mutations have been linked to RP.^1,2^ Gene therapy targeting specific mutations, such as *RPE65*, has been approved for treating RP patients.^3–5^ Given the high genetic diversity of RP, pursuing a mutation-independent therapeutic strategy to slow disease progression and preserve vision is a logical and promising approach.

Stem cell-based therapies offer promising avenues for treating retinal degenerative diseases through two main strategies: cell replacement and preservation of host cells. Replacement therapies, such as RPE cell transplants for age-related macular degeneration (AMD), have shown mixed outcomes in animal models and clinical trials.^6,7^ Similarly, photoreceptor transplant at the advanced stages of retinal degeneration have shown promising results in animal studies; however, establishing synaptic connections with host retinal cells remains challenging.^6,8,9^ Therapies that target preservation of host retinal cells have employed various stem cells in both preclinical and clinical settings, including retinal progenitor cells (RPCs)^10^, mesenchymal stem cells (MSCs)^11–13^, and human neural progenitor cells (hNPCs).^14–16^ Our preclinical studies demonstrate that hNPCs delivered to the subretinal space of animal model survive long-term, migrate extensively, and protect vision by providing trophic support, regulating immune response, and enhancing antioxidant defense mechanisms by upregulating Nrf2.^17–19^ These findings have led to the delivery of clinical-grade hNPCs (termed CNS10-NPC) to RP patients in an ongoing phase I/2a clinical trial (NCT04284293).

While stem cell-based therapies hold promise for retinal disease, long-term protection is compromised due to efficacy deterioration over time, as widely reported from both preclinical and clinical studies. Key questions remain about the behavior of grafted stem cells and host retina in a progressively degenerative environment, including how grafted cells adapt to this environment, how preserved photoreceptors evolve over time, and how grafted stem cells interact with host retinal cells.

Here, we investigated the changes in hNPCs and host retinal cells over time following subretinal injection into the Royal College of Surgeons (RCS) rat, a well-characterized model for retinal degeneration. Using single-cell RNA sequencing (scRNA-seq), along with histological and functional assessments, we analyzed the transcriptomic remodeling of hNPCs and host retinal cells and their interactions. Fourteen hNPC-subpopulations with distinct transcriptional states were identified, indicating dynamic changes in grafted cells over time. Top-regulated genes identified in grafted cells included mesencephalic astrocyte-derived neurotrophic factor (*MANF*), myeloid-derived growth factor (*MYDGF*), midkine (*MDK*), and pleiotrophin (*PTN*), which are newly identified trophic factors in hNPCs. The host retina showed temporal changes in phototransduction, visual perception, detection of light stimulus, glial reaction, and metabolic processes. Notably, the signaling interaction and strength between hNPCs and host retinal cells decreased over time. Grafted hNPCs protect vision through multiple modalities, including trophic factor release, extracellular matrix remodeling, metabolic activity regulation, and reduction of apoptosis, oxidative stress, and inflammation. The host retinal degenerative environment plays a critical role in affecting graft survival and function. These findings provide insight into the dynamic changes of grafted hNPCs and host retinal cells and emphasize grafting stem cells together with improving the host retinal environment is likely needed for long-term vision protection.

## RESULTS

### Temporal transcriptomic changes in grafted hNPCs within the degenerative retinal environment

To study the molecular insights of therapeutic effect and gradual changes of the grafted hNPCs over time in the degenerative retinal environment, hNPCs were delivered to the subretinal space of RCS rats at postnatal day (P) 21, followed by visual function evaluation at P60 and P90 (**Figure S1A**). Visual acuity and ERG b-wave amplitude were significantly higher for hNPC-treated eyes compared to untreated at P60 and P90 (**Figure S1B-C**). Notably, while visual acuity remained stable, ERG b-wave amplitudes decreased over time in hNPC-treated eyes. Histological evaluation of retinal sections also revealed a reduction in the photoreceptor nuclei in the outer nuclear layer of hNPC-treated retinas at P90 (6–8 layers) compared to P60 (8–10 layers; **Figure S1D-G**).

To understand the molecular underpinnings of these changes over time at single-cell resolution, the grafted-hNPCs and host retinal cells were collected at P60 and P90 and processed for scRNA sequencing (**Figure S1A**). High-quality cells (*in vitro*: 5502; *in vivo* P60: 4943; *in vivo* P90: 5536, **Figure S2A-D**) were collected by *in silico* sorting from the merged scRNA seq datasets of all time points (**Figure S2E-G**). Next, we identify the distinct cell-type clusters using known human and retinal cell-specific marker genes (**Figure S2H**). Six clusters were present for hNPCs, photoreceptors (PR), Müller glia, retinal ganglion cells (RGCs), bipolar cells, and microglia (**Figure 1A**). Subsequently, we subset 8,088 high-quality hNPCs (*in vitro* hNPCs: 5502; *in vivo* P60 hNPCs: 1575; *in vivo* P90 hNPCs: 1011) and performed unsupervised clustering on the batch-corrected, integrated transcriptomic dataset, resulting in 14 distinct subpopulations (**Figure 1B**), which were profiled and annotated based on the expression of well-established marker genes (**Figure 1C and S2I-J**).

**Figure 1:**
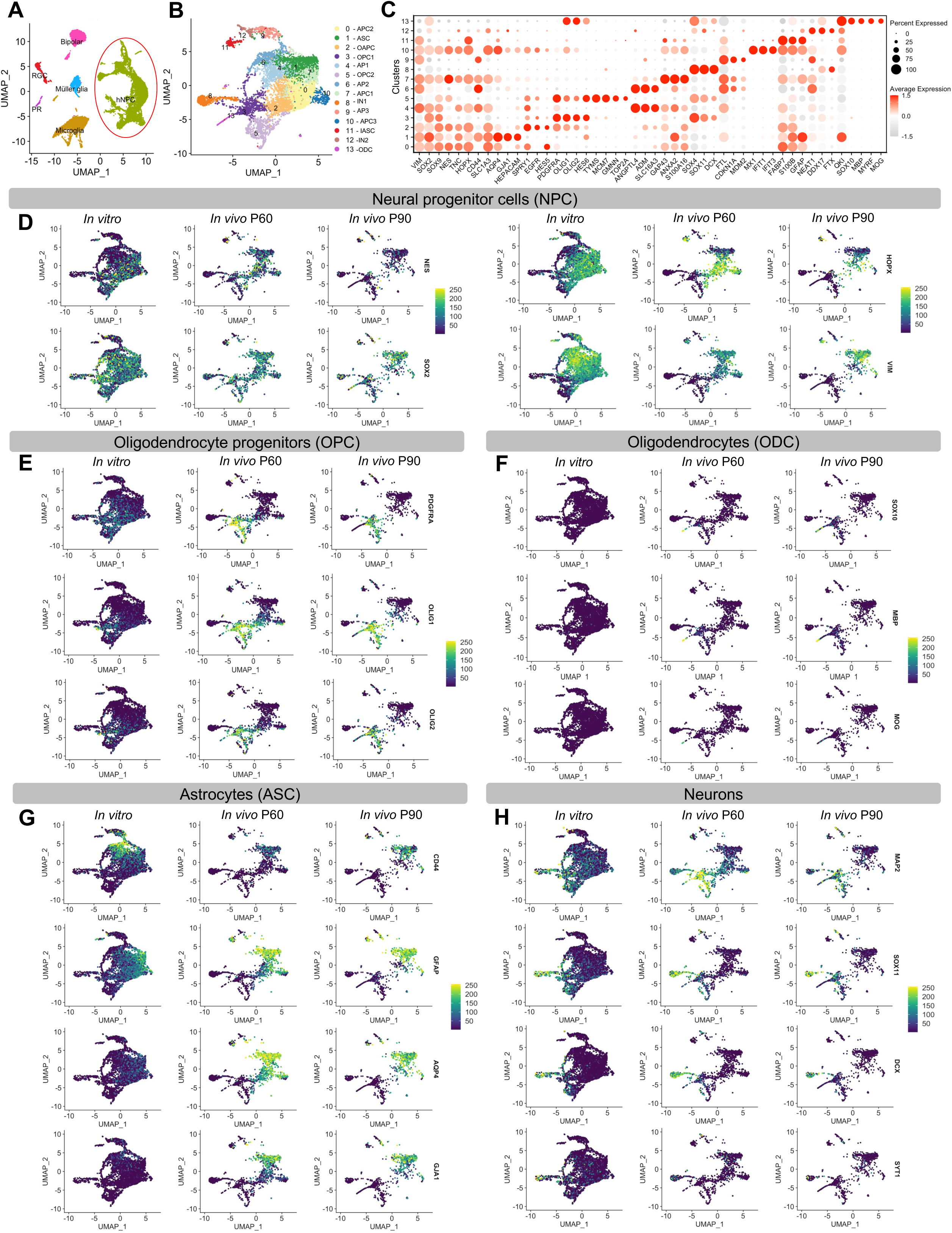
Single-cell transcriptomic analysis of subretinally grafted hNPCs in the RCS retina over time. **(A)** Uniform manifold approximation and projection (UMAP) plot of 15,981 cells profiled with single-cell RNA-seq (scRNA seq) showing grafted hNPCs and host rat retinal cell types. **(B)** UMAP plot of 8,088 hNPCs showing graph-based clustering of different hNPC-subpopulations, color-coded by cell type-specific annotated identities based on expressed transcripts of known genes. ASC, astrocytes; APC, astrocyte progenitor cells; AP, astrocyte precursor; IN, immature neurons; InN, interneurons; OAPC, oligodendrocyte astrocyte progenitor cells; ODC, oligodendrocytes; OPC, oligodendrocyte precursor cells, hNPC, human neural progenitor cells. **(C)** Dot plot showing the average expression of cell type-specific marker genes for transcriptomically defined clusters from scRNA seq. **(D-H)** UMAP plots of *in vitro* hNPCs and subretinally transplanted *in vivo* hNPCs, showing the expression patterns of cell type-specific marker genes of NPC **(D)**, OPC **(E)**, ODC **(F)**, ASC **(G)**, and neurons **(H)**.

The 14 transcriptomically distinct hNPC subpopulations were classified as *SOX9^+^/NES^+^/HOPX/SLC1A3^+^* astrocyte progenitor cells (APC; Clusters 0, n=1316; 7, n= 531, and 10, n=224), *CD44^+^/AQP4^+^/GJA1^+^/FLT^+^*astrocyte precursors (APs; Clusters 4, n=711; 6 n=533; and 9, n=337), *GFAP^+^/S100B^+^/FABP7^+^*immature astrocytes (IASC; Cluster 11, n=185), *GFAP^+^/AQP4^+^/GJA1^+^*mature astrocytes (ASC; Cluster 1 n=1042), *PDGFRA^+^/OLIG1^+^/OLIG2^+/^SLC1A3^+^/S100B^+^*oligodendrocyte astrocyte progenitor cells (OAPCs; Cluster 2; n=1042), *PDGFRA^+^/OLIG1^+^/OLIG2^+^*oligodendrocyte precursor cells (OPC; Clusters 3, n=755; and 5, n=708), small subpopulation of *SOX10^+^/MBP^+^/MYRF^+^* oligodendrocytes (OL; Cluster:13; n=114), *DCX^+^/SOX4^+^/SOX11*^+^ immature neurons (IN1; Cluster: 8; n=505) and *DDX17^+^/FTX^+^*immature neurons (IN2; Cluster: 12; n=166) (**Figure 1C and S2K-L**).

To explore the impact of the host degenerative retinal environment on the grafted hNPCs over time, the cluster-specific gene expression profile of hNPCs both before grafting and at P60 and P90 after transplantation was compared (**Figure 1D-H and S3A-F**). We found that clusters 0, 2, 5, 7, and 10 of grafted hNPCs expressed progenitor marker genes (*NES, SOX2*) over time (**Figure 1D**). We also observed that grafted hNPC clusters associated with astrocyte lineage (clusters: 0, 1, 7, 10, and 11) expressed mature astrocyte marker genes (*AQP4, GJA1*) compared to *in vitro* hNPCs with astrocyte lineage, suggesting prevalence of mature astrocyte states of grafted hNPCs *in vivo* (**Figure 1G**). Interestingly, we also noticed that cluster 2 of grafted hNPCs are enriched for both astrocyte-(*S100B, GFAP*) and oligodendrocyte-specific (*PDGFRA, OLIG1, OLIG2*) genes (**Figure S3C**). Further, we found that clusters 3 and 5 of grafted hNPCs are enriched for oligodendrocyte precursor markers (*PDGERA, OLIG1, OLIG2)* compared to clusters 3 and 5 of hNPCs *in vitro,* suggesting the acquisition of oligodendrocyte fate of grafted hNPCs in the host degenerative retinal environment (**Figure 1E and S3B**). Like previous studies^20–22^, both clusters 3 and 5 (oligodendrocyte precursor cell clusters: OPC1 and 2) also showed the enrichment of genes involved in neuron differentiation and development (*HES6, SOX11, DCX*; **Figure S3D-E**), indicating that they may have the capability to differentiate into neurons *in vivo*. Oligodendrocyte fate acquisition of grafted hNPCs is further supported by the presence of a small oligodendrocyte cluster enriched with mature oligodendrocyte genes (*MBP, MOB*), identified as cluster 13 only in grafted hNPCs, but absent in the *in vitro* hNPCs (**Figure 1F**). Gene ontology (GO) term enrichment analysis of upregulated genes in cluster 13 also revealed the enrichment of biological processes related to oligodendrocyte differentiation, myelination, and galactolipid biosynthesis (**Figure S3F**). Clusters 8 and 12 were enriched with early-developing neuronal genes (Cluster 8: *SOX4, SOX11, DCX*; and Cluster 12: *SOX11, DDX17)* in grafted hNPCs compared to *in vitro* hNPCs (Figure 1C, H **and S3B**). Clusters 4, 6, and 9 which are enriched with marker genes of different astrocyte maturation stages (*CD44, HEPACAM, ALDOC, SLC16A3*) were absent in grafted hNPCs (**Figure 1C and S3A-B**). Overall, these findings indicate that the majority of hNPCs acquire astrocytic states over time within the host retinal environment.

### Role of neurotrophic factors in enhancing hNPCs survival and retinal protection

We have previously shown that hNPCs provide retinal protection by secreting various neurotrophic factors, including insulin-like growth factor-1 (IGF-1) and fibroblast growth factor-2 (FGF-2)^14^. The decline in therapeutic efficacy over time may be partially due to reduced secretion of neurotrophic factors. Here, we found that in addition to *FGF*, the hNPCs expressed mesencephalic astrocyte-derived neurotrophic factor (*MANF*), Myeloid-derived growth factor (*MYDGF),* midkine (*MDK*) and pleiotrophin *(PTN)* both *in vitro* and *in vivo* hNPCs (**Figure 2 and S4**) as well as the expression of their receptors (**Figure 2A-C**). Notably, *MANF, MYDGF, MDK,* and *PTN* as well as their receptors were expressed at relatively high levels across the subpopulations of hNPCs (**Figure S4**). Interestingly, there was a significantly higher expression in *MDK*, *PTN*, and their receptors (*PTPRZ1, CSPG5)* in P60 compared to P90 grafted hNPCs (**Figure 2A**). MDK and PTN have been shown to promote the differentiation, migration, and survival of neural stem and progenitor cells.^23–27^ These trophic factors provide neuroprotection, reduce oxidative stress, and have anti-apoptotic effects.^23,25,28^ The autocrine action of *MDK* and *PTN*, via their receptors on hNPCs, promotes their stability, survival, and migration in the degenerative retinal environment. In addition, secreted MDK and PTN might act paracrinally on Müller glia through their receptors and protect photoreceptors^29^. We found a significant increase in *MANF, MYDGF*, and their receptor *KDELR1* in grafted hNPCs at P90 compared to P60 (**Figure 2B**). MANF has significant neuro-regenerative potential and protects photoreceptors from degeneration.^30–32^ MYDGF promotes NPC survival by maintaining their stemness.^33,34^ Both MANF and MYDGF can inhibit apoptosis, oxidative stress, inflammation, and enhance metabolic activity.^33,35–38^ Additionally, they can exhibit both autocrine and paracrine effects. Increased expression of *MANF*, *MYDGF,* and their receptors in grafted hNPCs at P90 compared to P60, suggest their autocrine neuroprotection to hNPCs, supporting their self-protection, stability, and survival in the progressive degenerative retinal environment. Moreover, increased secretory *MANF* and *MYDGF*, alongside their autocrine protection to hNPCs, may also protect photoreceptors via KDELR through paracrine action.

**Figure 2:**
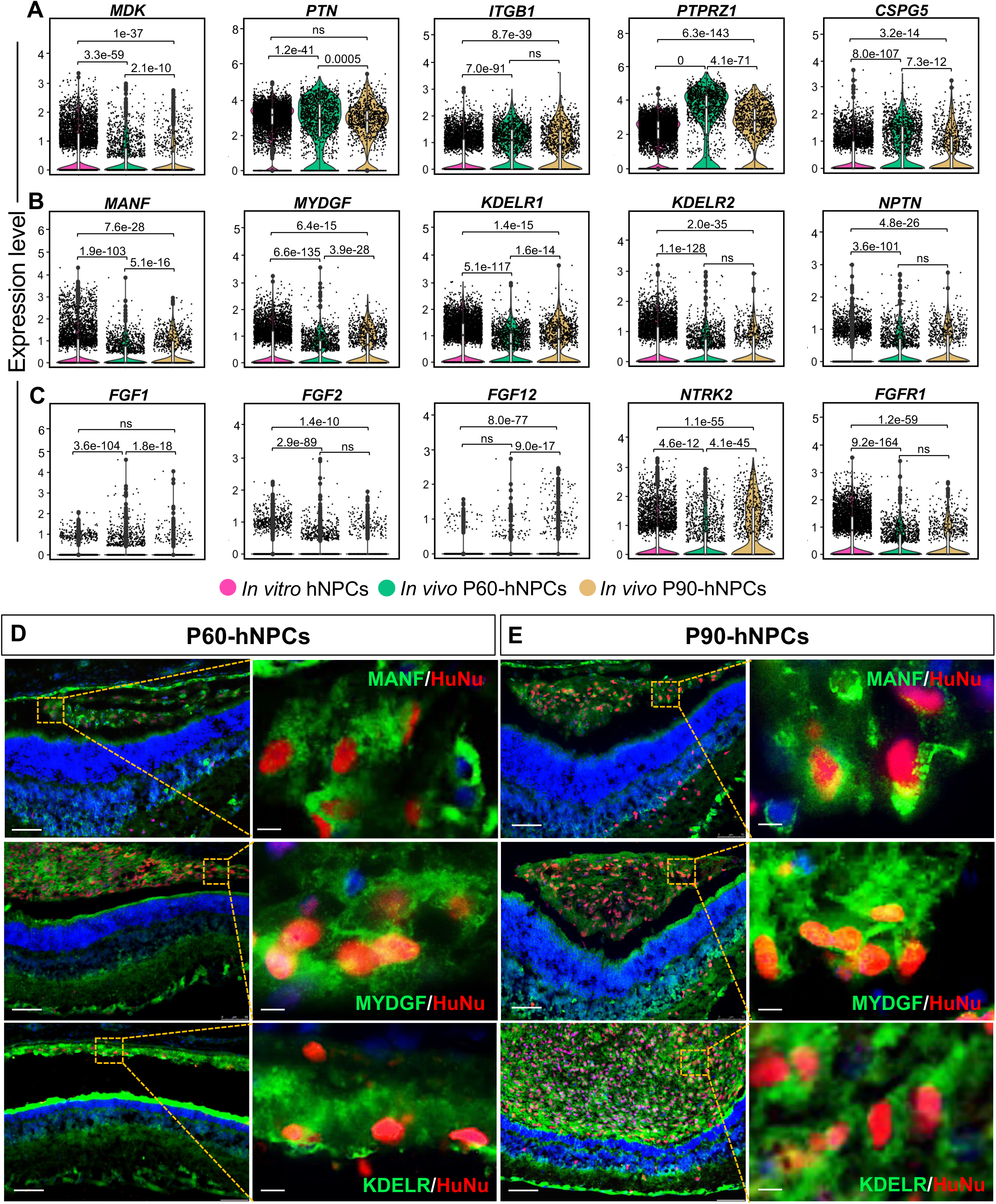
Identification of trophic and growth factors and their receptors in hNPCs over time. **(A)** Violin plots showing the expression of major growth factors *MDK* and *PTN* and their receptors (*ITGB1, PTPRZ1* and *CSPG5*) in *in vitro* and grafted P60 and P90 hNPCs. **(B)** Violin plots showing the expression of novel neurotrophic and growth factor *MANF* and *MYDGF* and their receptors (KDELR1, 2 and NPTN) in *in vitro* and grafted P60 and P90 hNPCs. **(C)** Violin plots showing the expression of neurotrophic factors (*FGFs*) and their receptors (*NTRK2* and *FGFR1*) in *in vitro* and grafted P60 and P90 hNPCs. **(D-E)** Confocal Immunofluorescence (IF) images from hNPCs at P60 and P90 time points after subretinally grafting at P21 in the RCS retina, showing MANF (green, upper panel) and MYDGF (green, middle panel) stained with a human-specific nuclear marker (HuNu: red). Double immunostaining of KDELR (green, lower panel) and human nuclear marker (HuNu, red) showing KDELR in P90 hNPCs. Scale bars, 50μm and 10μm.

We validated our findings using immunofluorescence of MANF, MYDGF, and their receptor KDELR in grafted hNPCs at P60 and P90. Double immunolabeling using human-specific nuclear marker HuNu (red) and MANF (green) demonstrated cytoplasmic and perinuclear puncta staining of MANF at both P60 and P90 (**Figure 2D-E**, upper panel). Further, double immunofluorescence using HuNu (red) and MYDGF (green) demonstrated intense cytoplasmic, perinuclear, and nuclear staining of MYDGF in grafted hNPCs (**Figure 2D-E**, middle panel). Next, double immunofluorescence using HuNu (red) and KDELR (green) demonstrated intense cytoplasmic staining of KDELR in hNPCs (**Figure 2D-E**, lower panel). Altogether, these findings suggest that *MDK* and *PTN* may promote grafted hNPC migration and survival through their autocrine and paracrine action; *MANF* and *MYDGF* promote grafted hNPC survival and function via inhibiting cellular oxidative stress, inflammation, and immune response as degeneration progresses.

### Biological adaptation and functional shifts in grafted hNPCs in a degenerative retinal environment

Grafted hNPCs undergo extensive biological changes over time in response to the host degenerative retinal environment. To investigate the adaptation and functional shifts, we compared transcriptomic profiles of grafted hNPCs at P60 and P90 with *in vitro* hNPCs (**Figure 3A-B**). At P60, grafted hNPCs displayed increased expression of genes involved in differentiation, neuron projection morphogenesis, synaptic signaling, and survival pathways, reflecting their integration and functional maturation within the host retina. By P90, fewer genes showed differential expression, indicating stabilization of hNPCs in the host environment (**Figure 3A-D**). Common pathways enriched across P60 and P90, including cellular processes supporting differentiation, migration, axonogenesis, endocytosis, and autophagy, as well as suppression of immune activation (e.g., TGF-β regulation). In contrast, biological processes related to neurotrophin and mTOR signaling, negative regulation of apoptosis, protein processing in ER, ErbB signaling, and metabolic pathways, including glycolysis and pyruvate metabolism, were consistently downregulated, highlighting a shift in energy utilization and metabolic activity over time (**Figure 3E**).

**Figure 3:**
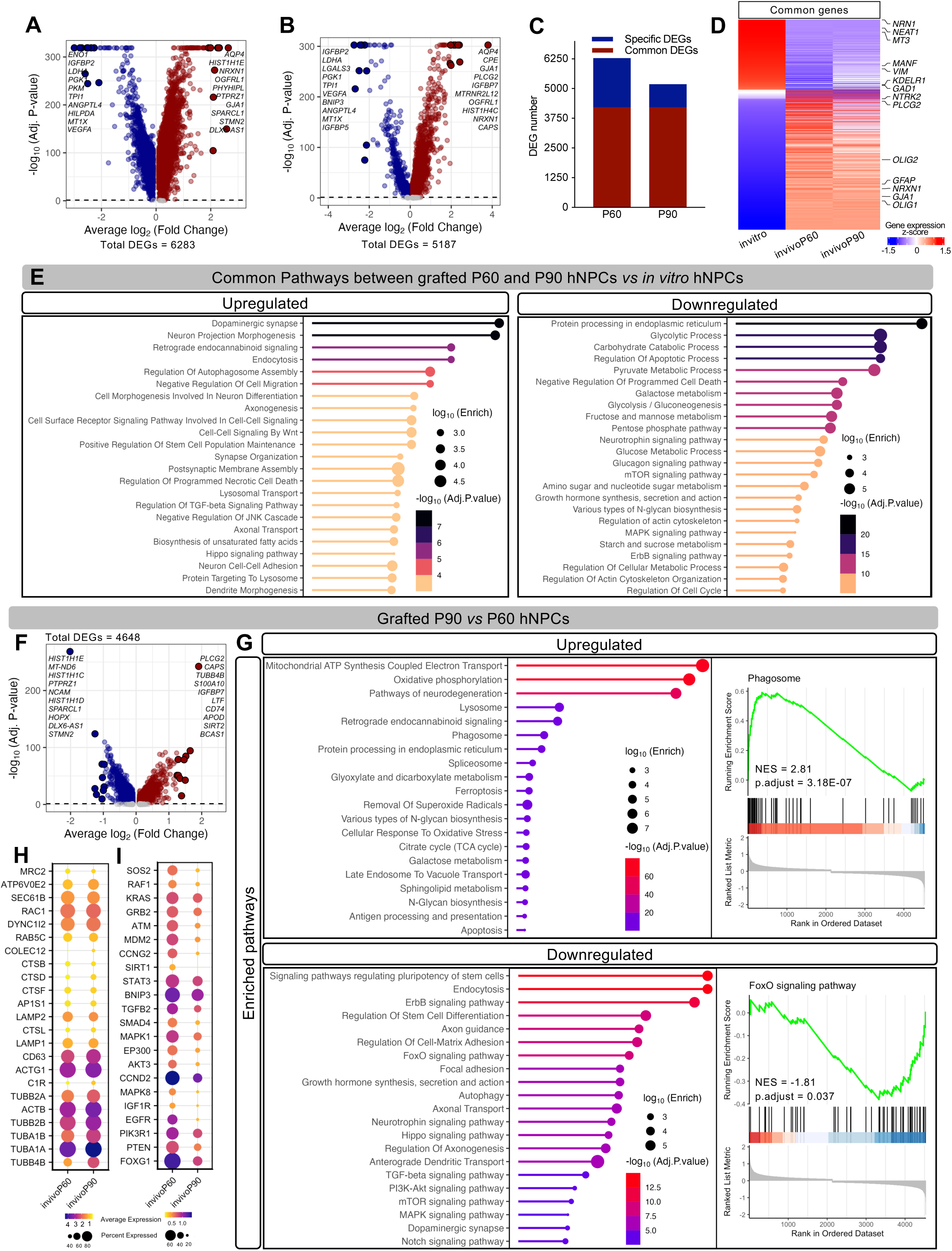
Transcriptomic remodeling of grafted hNPCs in RCS retina over time. **(A-B)** Volcano plots of differentially expressed genes (DEGs) showing the effect size (average log2FC) and significance levels (Adj. P-values) from grafted P60 **(A)** and P90 **(B)** hNPCs versus *in vitro* hNPCs respectively. The top 10 up- and down-regulated significant genes by effect size are labeled. The maroon are upregulated genes, dark blue are downregulated genes, and gray are non-significant. **(C)** Bar graph showing the number of common and specific DEGs between grafted P60 and P90-hNPCs versus *in vitro* hNPCs. **(D)** Heatmap showing the common DEGs among grafted P60 and P90-hNPCs versus *in vitro* hNPCs. **(E)** Lollipop plots showing the selected pathway enrichment results from DEGs that were commonly upregulated and downregulated in both grafted P60 and P90-hNPCs compared to *in vitro* hNPCs. **(F)** Volcano plot of DEGs showing the effect size and significance level from *in vivo* P90 versus P60-hNPCs. The top 10 up- and down-regulated significant genes by effect size are labeled. The maroon are upregulated genes, dark blue are downregulated genes, and gray are non-significant. **(G)** Lollipop plots showing the selected pathway enrichment results from DEGs that were upregulated and downregulated in grafted P90 compared to P60-hNPCs. **(H-I)** Dot plots showing the expression of phagocytic and lysosomal genes **(H)** and FoxO signaling genes **(I)** in grafted P60 and P90-hNPCs.

Temporal analyses revealed distinct transcriptomic changes in P60 and P90 hNPCs. At P60, grafted hNPCs upregulated genes linked to neuron development, adhesion, and axonogenesis while downregulated genes related to metabolic pathways, reflecting early adaptation to the host retina (**Figure S5A-H**). By P90, there was increased expression of genes associated with glial differentiation and immune responses, along with further suppression of metabolic pathways, suggesting a shift toward immune and stress-responsive phenotypes as degeneration progressed (**Figure S5I-O**). These changes reflect ongoing adaptation and maturation of hNPCs within a progressively degenerative environment.

To understand the impact of the degenerative retinal environment on the transcriptomic remodeling of grafted hNPCs over time, we conducted a comparative transcriptomic analysis between P60 and P90 hNPCs (**Figure 3F**). At P90, hNPCs exhibited enhanced expression of genes related to phagocytosis, energy metabolism (e.g., mitochondrial ATP synthesis coupled electron transport, oxidative phosphorylation), and cellular stress and death (e.g., response to oxidative stress, apoptosis) (**Figure 3G**), reflecting increased metabolic demands and stress responses. Notably, the upregulation of lysosomal and phagosome-related genes (e.g., *CTSD*, *LAMP1*, *LAMP2*) suggests active clearance of photoreceptor outer segments (POS) from the subretinal space, a protective function that may aid photoreceptor survival (**Figure 3H**).^19,39^ However, the downregulation of pathways such as FoxO signaling, TGF-β signaling, and neurotrophin signaling, coupled with increased oxidative stress and immune activation, indicates a gradual shift in hNPCs toward a stress-responsive phenotype as degeneration progresses (**Figure 3G-I, S6A-L**), characterized by reduced trophic support and survival capacity.

### Transcriptomic remodeling of retinal cells following hNPC transplantation

To assess the impact of hNPC treatment on the host degenerative retina, we performed scRNA seq on retinal cells from untreated and hNPC-treated retinas at P60 and P90, alongside wild-type (WT) controls.^40^ A total of 118,478 high-quality cells, encompassing all major retinal cell types, were analyzed (**Figure S7A-G**). These included rods (14,675), cones (4,341), rod bipolar cells (RBCs: 40,882), cone bipolar cells (CBCs: 9,650 cells), Müller glia (MULG: 14,614), retinal ganglion cells (RGCs: 9,654), amacrine cells (AC: 2,577), as well as microglia (MGs: 15,084), macrophages (MO: 519), and endothelial cells (ECs: 6,482) (**Figure S7H-I**). Retinal degeneration in RP marked by progressive loss of rods and cones, with corresponding changes in Müller glia and microglia. Müller glia, known to support photoreceptor health through neurotrophic and metabolic functions, exhibited altered transcriptional profiles, reflecting their response to retinal stress.^41^ Similarly, microglia showed signs of activation and infiltration, consistent with their role in degenerative processes.^42^ Hence, we focused on rods, cones, Müller glia, and microglia for further downstream analysis. Our analysis demonstrated that hNPC transplantation induced distinct transcriptomic changes in retinal cell types, preserving photoreceptor gene expression linked to visual perception and maintenance, upregulating neurotrophic pathways in Müller glia, and reducing pro-inflammatory gene expression in microglia. Overall, these findings demonstrate the capability of hNPCs in remodeling the retinal environment to support photoreceptor survival and function.

### Photoreceptor dysfunction and rescue mechanisms during retinal degeneration and hNPC-treatment

To understand the biological changes in photoreceptors during retinal degeneration, we analyzed transcriptional signatures in rods and cones at P60 and P90. In untreated retinas, 1,811 and 1,685 differentially expressed genes (DEGs) were identified in rods at P60 and P90, respectively, while cones exhibited 1,634 and 2,664 DEGs (**Figure 4A-C**). Gene ontology (GO) analysis revealed downregulation of photoreceptor signaling, phototransduction pathway, G protein transmit signals (*Gnb1, Cnga1, Pde6b*) and TOR signaling (*Ddit4, Lars1*) in rods at P60, alongside progressive declines in genes essential for visual perception, photoreceptor maintenance, and glucose metabolism by P90, compared to WT (**Figure 4D, F**). Concurrently, untreated rods showed upregulation of genes associated with synapse organization, axonogenesis (*Lrrtm4, Nrxn3, Gnb3),* and calcium channel activity (*Calm1, Cacna2d1, Robo2*), ATP synthesis, and neurotransmitter secretion/transport and downregulation of genes related to glycolysis (*Hk2, Ldha*), indicating metabolic dysfunction and neuronal remodeling (**Figure 4E and G**).

**Figure 4:**
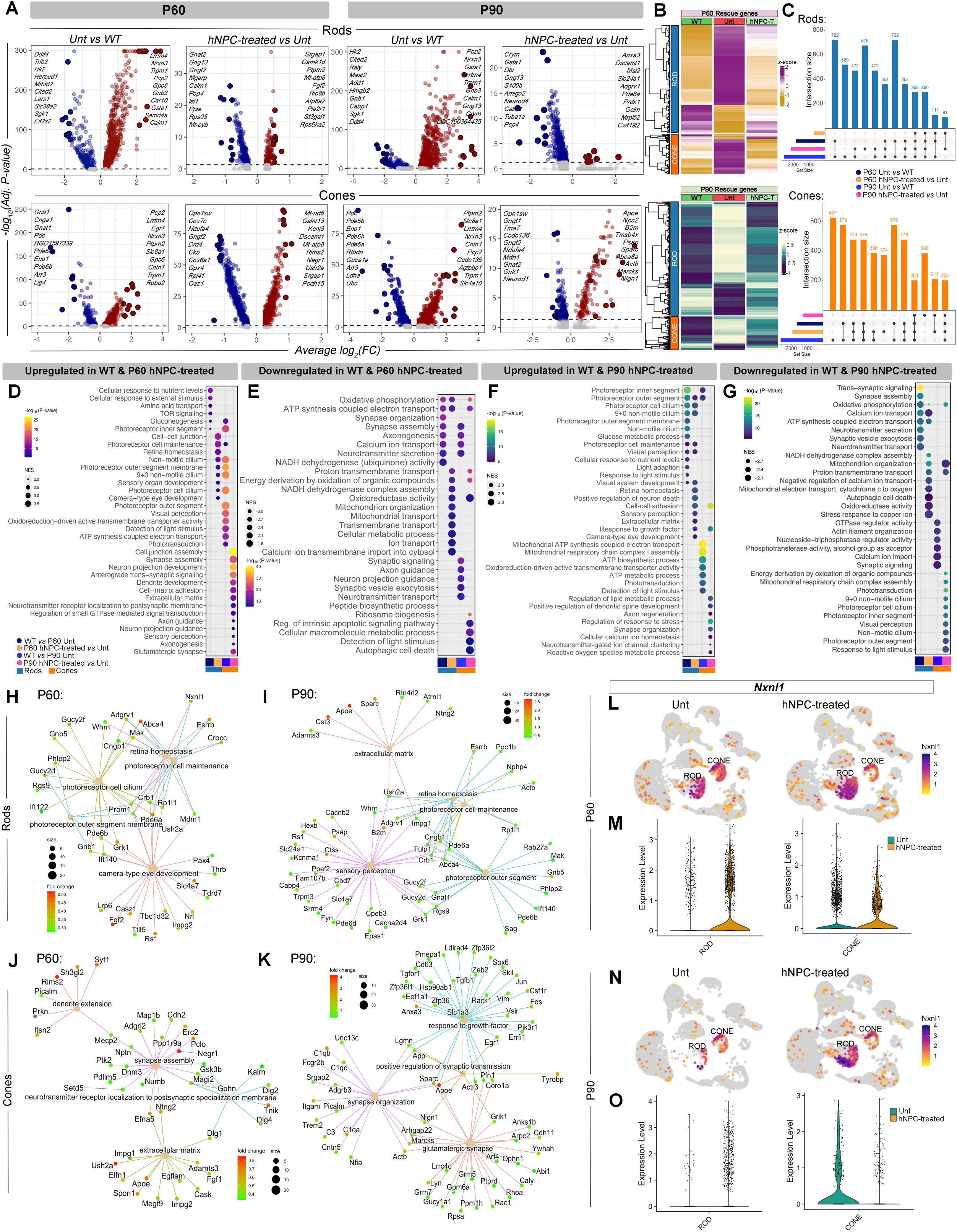
Transcriptomic remodeling in RCS rat retina highlights hNPC-induced protective changes in rods and cones over time. **(A)** Volcano plots of DEGs showing the effect sizes (average log2FC) and significance levels (Adj. P-values) from untreated (Unt) *versus* wild type (WT) and hNPC-treated *versus* untreated P60 and P90 rods and cones. The top 10 significantly up- and down-regulated genes by effect size are labeled. Maroon circles are upregulated, dark blue circles are downregulated, and gray circles are non-significant genes. **(B)** Heatmaps of P60 and P90 rods and cones showing differential expression of the rod and cone-specific ‘rescue’ (functionally protective) genes over time. **(C)** Upset plots showing the overlap between the sets of differentially expressed genes in P60 and P90 rods and cones among different groups. **(D-E)** Dot plot showing the selected pathway enrichment results from DEGs that were up- and down-regulated in wild type and P60 hNPC-treated rods and cones compared to P60 untreated. **(F-G)** Dot plot showing the selected pathway enrichment results from DEGs that were up- and down-regulated in wild type and P90 hNPC-treated rods and cones compared to P90 untreated. **(H-I)** Cnet plots showing the interactions of selected up-regulated pathways, and their associated genes in P60 and P90 hNPC-treated rods **(H and I)** and cones **(J and K)** compared to untreated. The size of each circle associated with each pathway represents the number of genes enriched in each pathway, and the color of each gene represents the log2FC (fold change). **(L-O)** Feature plots **(L and N)** and violin plots **(M and O)** showing the expression of *Nxnl1* in P60 and P90 hNPC-treated rods and cones compared to untreated.

In untreated retinas, cones displayed similar transcriptional trends, with reduced expression of genes related to phototransduction, visual perception, and ATP metabolism alongside increased expression of synaptic signaling and axon outgrowth genes (**Figure 4E and G**). These findings suggest rods and cones experience metabolic stress and compromised function during degeneration, with compensatory neuronal reorganization to maintain connectivity.

To elucidate the photoreceptor protective mechanism after hNPC treatment, we conducted transcriptomic analysis between hNPC-treated and untreated rods and cones at P60 and P90. In rods, 688 differentially expressed genes were identified at P60 and 2,130 at P90 following hNPC treatment (**Figure 4A**). Of these, 457 DEGs at P60 and 1,198 at P90 were shared between untreated/WT and hNPC-treated/untreated. Importantly, among these shared DEGs, only 382 at P60 and 985 at P90 in hNPC-treated rods showed expression patterns similar to WT rods, defining them as rod-specific rescue genes (**Figure 4B-C**). Further analysis of time point-specific DEGs revealed upregulation of photoreceptor maintenance, photoreceptor cilium, retinal homeostasis, and glycolysis (*Hk2, Ldha*) in P60 and P90 hNPC-treated rods. Additionally, P90 hNPC-treated rods showed upregulation of genes involved in photoreceptor inner/outer segments, visual and sensory perception (*Cnga1, Rho, Pde6b*), highlighting enhanced photoreceptor integrity and function (**Figure 4D, F, H and J**). These data strongly demonstrated that hNPCs protect rods and slow down the progression of retinal degeneration.

Next, we evaluated the transcriptomic changes in cones at P60 and P90 following hNPC treatment. We identified 1,912 DEGs at P60 and 1,254 DEGs at P90 in hNPC-treated cones compared to untreated (**Figure 4A**) Among these, 761 at P60 and 592 at P90 were shared between untreated/WT and hNPC-treated/untreated cones. Intriguingly, only 172 DEGs in P60 and 289 in P90 hNPC-treated cones displayed expression patterns similar to WT cones, annotated as cone-specific rescue genes (**Figure 4B-C**). GO analysis of time point-specific DEGs revealed that at P60, hNPC-treated cones exhibited increased expression of genes associated with anterograde trans-synaptic signaling and GTPase-mediated signal transduction (**Figure 4D and J**) while, at P90 hNPC-treated cones demonstrated upregulation of genes involved in synapse organization and lipid metabolism (**Figure 4F and K**). However, mitochondrial organization and transport genes were consistently downregulated across time points in treated rods and cones, indicating potential energy production deficits critical for sustained photoreceptor function (**Figure 4E, G, and S8A-B**). In contrast, compared to hNPC treatment, untreated P60 cones, and P90 rods showed upregulated genes associated with autophagic cell death, consistent with previous findings suggesting that dysregulated autophagy contributes to photoreceptor cell death in rodent models for retinal degeneration.^19,43,44^ Moreover, P60 untreated cones also showed upregulation of genes associated with light detection and oxidoreductase activity compared to hNPC-treated cones (**Figure 4E and S8C**). Notably, untreated P90 cones retained some functionality, with higher expression of phototransduction and light response genes compared to hNPC-treated cones, suggesting residual light responsiveness despite degeneration (**Figure 4F and S8D**) These results align with recent studies indicating that cones retain partial functionality even as degeneration progresses.^45^

hNPC treatment also significantly increased the expression of *Nxnl1*, encoding the rod-derived cone viability factor (RDCVF)^46–49^, which reduces oxidative stress^50^ and promotes aerobic glycolysis to support cone survival^51,52^, underscoring the therapeutic potential of hNPCS (**Figure 4l-O and S9**). Interestingly, untreated cones at P60 also exhibited elevated *Nxnl1* expression compared to hNPC-treated cones, potentially reflecting an intrinsic compensatory response to rod loss. However, downregulation of mitochondrial genes in hNPC-treated photoreceptors highlights a need to address energy deficits to enhance long-term photoreceptor survival.^53–55^ These findings demonstrate that hNPC treatment preserves retinal integrity by promoting photoreceptor function and maintenance despite limitations in sustaining mitochondrial energy production at advanced stage of degeneration.

### Dynamic transcriptomic reprogramming of Müller glia and microglia by hNPC transplantation

Our previous studies demonstrated suppression of Müller cell gliosis and microglial activation following hNPC transplantation in the RCS rat.^18,19^ However, how hNPCs modulate the transcriptomic signatures of Müller glia and microglia remains unknown. To explore these changes, we analyzed Müller glial and microglial transcriptomes at P60 and P90, revealing distinct temporal and functional reprogramming patterns (**Figure 5A-K**).

**Figure 5:**
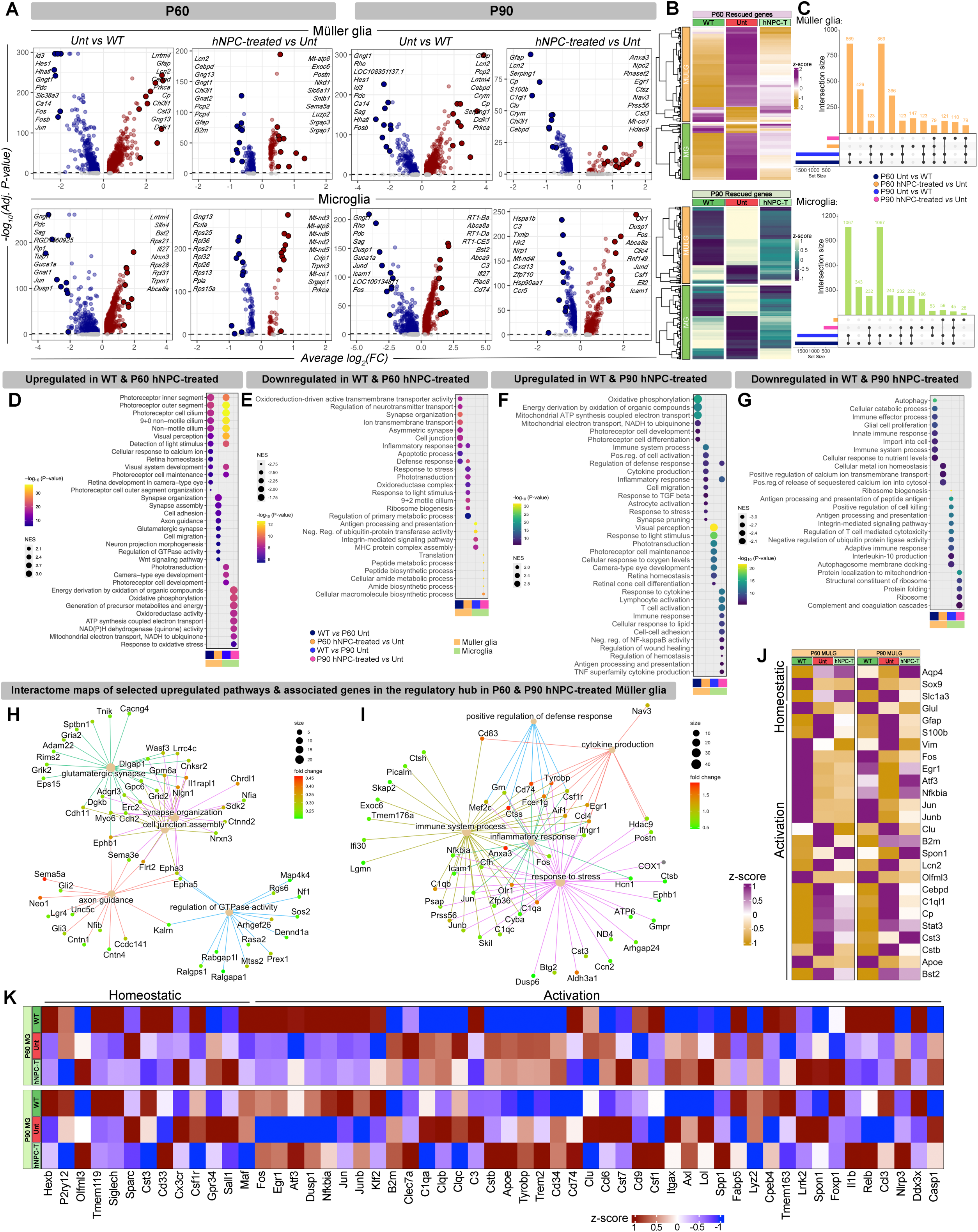
hNPC-induced immunomodulation *via* transcriptomic remodeling of Müller glia and Microglia. **(A)** Volcano plots of DEGs showing the effect sizes (average log2FC) and significance levels (Adj. P-values) from untreated (Unt) versus wild type (WT) and hNPC-treated versus untreated P60 and P90 Müller glia and microglia. The top 10 significantly up- and down-regulated genes by effect size are labeled. Maroon circles are upregulated, dark blue circles are downregulated, and gray circles are non-significant genes. **(B)** Heatmaps of P60 and P90 Müller glia and microglia showing differential expression of the Müller glia and microglia-specific ‘rescue’ (functionally protective) genes over time. **(C)** Upset plots showing the overlap between the sets of differentially expressed genes in P60 and P90 Müller glia and microglia among different groups. **(D-E)** Dot plot showing the selected pathway enrichment results from DEGs that were up- and down-regulated in wild type and P60 hNPC-treated Müller glia and microglia compared to P60 untreated. **(F-G)** Dot plot showing the selected pathway enrichment results from DEGs that were up- and down-regulated in wild type and P90 hNPC-treated Müller glia and microglia compared to P90 untreated. **(H-I)** Cnet plots showing the interactions of selected upregulated pathways, and their associated genes in P60 **(H)** and P90 **(I)** hNPC-treated Müller glia compared to untreated Müller glia. The size of each circle associated with each pathway represents the number of genes enriched in each pathway, and the color of each gene represents the log_2_ fold change (fold change). **(J)** Heatmaps demonstrating the expression of selected Müller glial homeostatic (non-reactive) and activation (reactive gliosis) genes in P60 and P90 wild-type (WT), untreated (Unt) and hNPC-treated (hNPCT) groups. **(K)** Heatmaps demonstrating the expression of selected microglial homeostatic and activation (early and late disease-associated) genes in P60 and P90 wild-type (WT), untreated (Unt) and hNPC treated (hNPCT) groups.

By comparing untreated Müller glia with WT, we first investigated the disease-associated changes in these cells over time. We identified 1,710 differentially expressed genes at P60 and 1,685 at P90 (**Figure 5A**). Untreated Müller glia displayed increased expression of genes associated with reactive gliosis (*Gfap, Lcn2*) at both early (P60) and late (P90) stages of the disease, compared to WT, concordant to other retinal degenerative diseases (**Figure 5D-G**).^56^ Specifically, at P60, untreated Müller glia exhibited downregulation of early responsive genes (*Egr1, Fos, Jun*) and upregulation of genes involved in inflammation, defense response, and apoptosis (*Ifi27, Cp, Cst3, Clu*), indicating the activation of stress and immune responses at the early stage of retinal degeneration. Additionally, at P60, untreated Müller glia also showed increased expression of genes (*Nlgn1, Ephb1, Nrxn3*) related to neurotransmitter transport and synapse organization, suggesting their compensatory role in remodeling and stabilizing synapses to support retinal connections at early-stage of disease. However, downregulation of genes involved in visual perception, photoreceptor cell maintenance, cellular response to calcium ions, and retina homeostasis reflects impaired support for photoreceptor function at the early stage. As the disease progresses, untreated Müller glia exhibited further upregulation of genes associated with glial cell proliferation (*Vim, Gfap, Clu*), innate immune response (*Apoe, C1s, Lcn2*), and autophagy (*Gabarap, Ctsd, Sqstm1*), coupled with downregulation of genes related to oxidative phosphorylation and mitochondrial ATP synthesis coupled electron transport, indicating metabolic dysfunction at late stage of disease. Consistent with previous studies, these findings suggest a shift in the Müller glial reactive state, transitioning from an early-stage compensatory to a late-stage decompensatory role with retinal degeneration progression.^57,58^

Differential gene expression analysis identified 1,801 DEGs at P60 and 1,707 DEGs at P90 in untreated microglia compared to WT (**Figure 5A**). At both time points, untreated microglia exhibited upregulation of genes involved in antigen processing and presentation (*B2m, Cd74*) and integrin-mediated signaling (*Tyrobp, Itgb1*). At P90, additional upregulation of genes related to the adaptive immune response (*B2m, C3*), interleukin-10 production (*Plcg2, Cd84*), and autophagosome membrane docking (*Vamp8, Calm1*) was observed (**Figure 5A**). However, genes associated with phototransduction, visual perception, photoreceptor cell maintenance, and retina homeostasis were downregulated (**Figure 5D and F**). Together, these findings suggest a dynamic transition in the microglial reactive state to mitigate cellular stress by enhancing adaptive immune response and reducing inflammation through secreting anti-inflammatory cytokines along with disease progression.

Comparative analysis between hNPC-treated and untreated Müller glia identified 478 DEGs at P60 and 461 DEGs at P90 (**Figure 5A**). Among these, 265 DEGs at P60 and 299 at P90 were shared across untreated/WT and hNPC-treated/untreated groups, with 159 (P60) and 261 (P90) displaying expression patterns similar to WT. These were classified as Müller glia-specific rescue genes (**Figure 5B-C**). GO analysis revealed that hNPC treatment at P60, upregulated genes associated with synapse organization (*Nlgn1, Nrxn3*), cell adhesion (*Sema5a, Neo1*), axon guidance (*Ephb1, Epha5*), and glutaminergic synapse (*Nlgn1, Gria2*), highlighting a neuroprotective role in maintaining synaptic stability and retinal integrity (**Figure 5D and H**). By P90, upregulated genes were linked to cytokine production (*Egr1, Csf1r*), cell migration (*Ccl4, Jun*), as well as immune, inflammatory, and defense responses (*C1qa, Nfkbia*), indicating adaptation to the progressively degenerative environment through protective immune functions (**Figure 5F and I**). Interestingly, at P60, hNPC-treated Müller glia exhibited downregulation of genes related to phototransduction and light response, while at P90, genes related to cellular metal ion homeostasis and calcium ion transport (*Mt2A, Smdt1, Calm1*) were downregulated (**Figure 5E and G**). However, key reactive gliosis genes (*Gfap, Vim*) were consistently downregulated in hNPC-treated Müller glia at both time points, indicating suppression of reactive gliosis following hNPC treatment (**Figure 5J**).

At P60, hNPC-treated microglia showed increased expression of genes related to oxidative phosphorylation, ATP synthesis, and NAD(P)H dehydrogenase (quinone) activity (*Mt-nd3, Mt-atp8*) (**Figure 5D**), reflecting enhanced homeostatic functions such as surveillance and phagocytosis, which rely on oxidative metabolism.^59^ By contrast, in inflammatory microglia, metabolic reprogramming suppresses their oxidative phosphorylation.^60^ However, by P90, hNPC-treated microglia showed upregulation of disease-associated microglial (DAM) genes^61^ related to inflammatory and immune responses (*Icam1, Clec7a, Apoe, Il1b, Spp1*) (**Figure 5F and K**) along with genes involved in the homeostatic process (*Igf1, Nfe2l*), lipid metabolism (*Apoe, Abca1, Trem2*), and negative regulation of TNF production (*Cd274, Tnfaip3*), suggesting an adaptive response to stabilize retinal homeostasis, modulate inflammation, and mitigate damage. Conversely, hNPC-treated microglia at P60 showed downregulation of genes involved in peptide metabolism (*Rpl30, Rps12*), while at P90, genes related to ribosome biogenesis (*Rplp1, Rps2*), protein folding (*Hsp90aa1, Hsph1*), and complement cascades (*C1qa, C3*) were downregulated. These changes reflect metabolic and functional reprogramming in microglia in response to degenerative retinal environment with disease progression.

These findings suggest that hNPC treatment dynamically modulates Müller glial and microglial responses over time to counteract the degenerative retinal environment. At P60, hNPCs promote neuroprotection by stabilizing synaptic connections,^19,62^ suppressing reactive gliosis, and enhancing microglial energy metabolism to support their surveillance and phagocytosis, critical for maintaining retinal homeostasis.^63,64^ By P90, Müller glia exhibit increased immune and inflammatory responses, lipid metabolism, and cytokine production. These results suggest a dynamic transcriptomic reprogramming, resulting in a neuroprotective macroglial phenotype, fueling protective phagocytosis and mitigating further retinal degeneration.^65,66^ In sum, hNPC treatment provides neuroprotection over time *via* dynamic transcriptomic reprogramming of glial cells within the progressive degenerative retinal environment.

### Temporal transcriptomic shifts in retinal cells highlight neuroprotective and adaptive responses to hNPC treatment

To explore how hNPC-treated retinal cells change their transcriptomes and assess their functional stability in a progressive degenerative retinal environment over time, we compared the transcriptomic profiles of P60 with P90 hNPC-treated rods, cones, Müller glia, and microglia. We found a total of 890 DEGs in rods, 1,075 in cones, 691 in Müller glia, and 757 in microglia (**Figure 6A-B**). Gene set overlap analysis demonstrated that up- and down-regulated DEGs with the largest effect sizes were shared across these cells (**Figure 6C-D**). At P60, rods and cones showed increased expression of genes related to photoreceptor inner/outer segments (*Gngt1, Cnga1, Rho*), phototransduction (*Rcvrn, Pde6g*), and visual perception (*Pdc, Guca1a, Rgs9, Cngb3*) and decreased expression of apoptosis (*Selenos, Btg2*) and phagocytosis-related genes (*Cst3, Ctsd*) (**Figure 6E-G and S10A-B**). Additionally, at P60, hNPC-treated cones also displayed upregulation of genes essential for photoreceptor maintenance (*Crb1, Cep290*), cone development (*Gnat2, Thrb*), mitochondrial bioenergetics pathways [NADH dehydrogenase (quinone) activity and ATP synthesis coupled electron transport (*Mt-nd6, Mt-nd5*)] (**Figure 6E and H**), indicating higher metabolic activity, important for cone function and maintenance. Conversely, at P90, cones showed increased expression of genes related to ROS metabolic processes (*Nfe2l2/Nrf2, Cyba*), while rods upregulated genes linked to calcium ion homeostasis (*Kcnma1, Calm1*), growth factor responses (*Nfkb1, Egr1, Fos*), and synapse formation (*Kif1a, Nrxn3*), indicative of adaptive responses to oxidative and metabolic stress in the progressive degenerative retinal environment.

**Figure 6:**
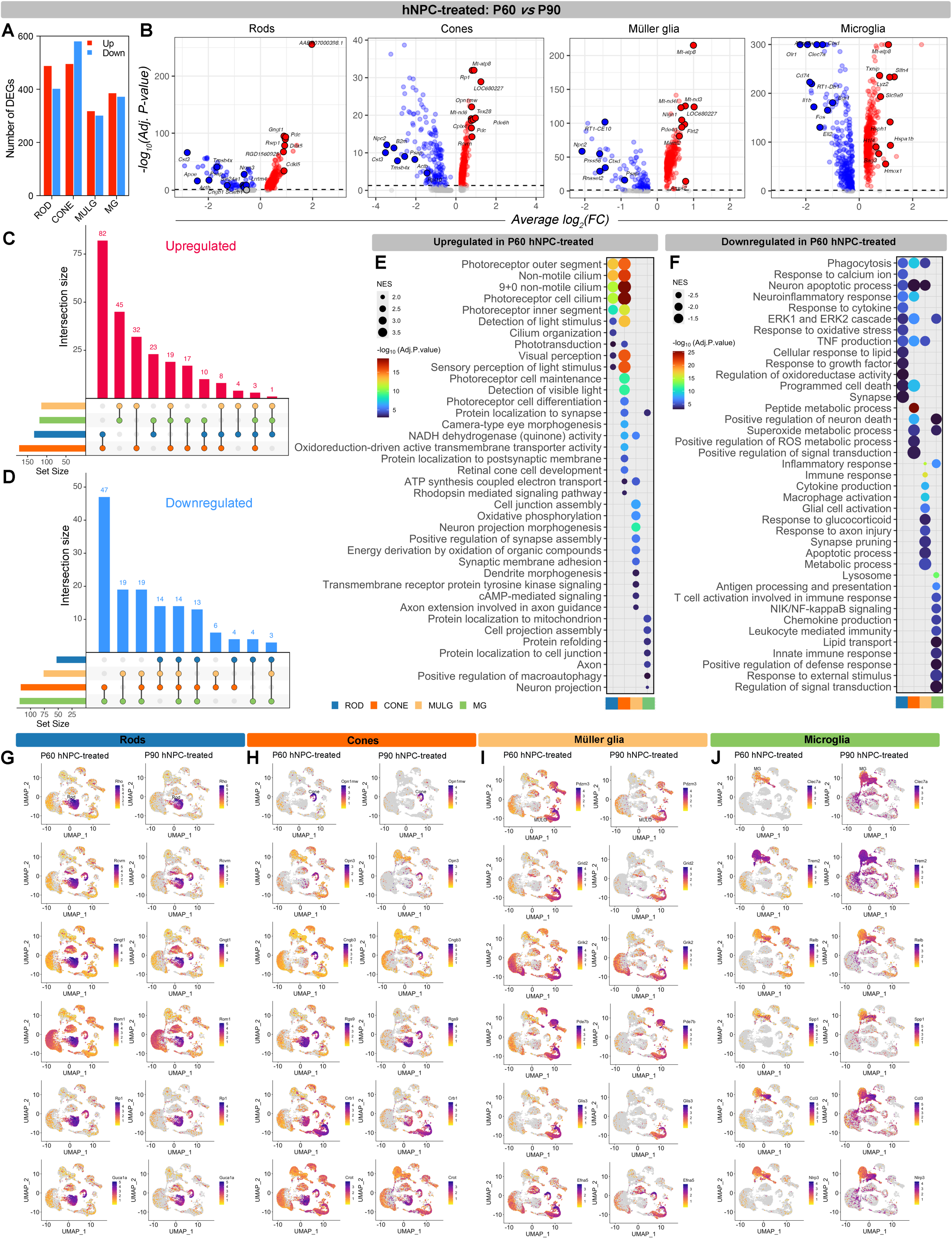
hNPC-induced temporal transcriptomic remodeling and compensatory responses maintain stability and function of protected photoreceptors. **(A)** Bar plots showing the number of differentially expressed up- and down-regulated genes between P60 and P90 hNPC-treated rods, cones, Müller glia and microglia. **(B)** Volcano plots of DEGs showing the effect sizes (average log2FC) and significance levels (Adj. P-values) from P60 versus P90 hNPC-treated rods, cones, Müller glia and microglia respectively. The top significantly up- and down-regulated genes by effect size are labeled. Red circles are upregulated, blue circles are downregulated, and gray circles are non-significant genes. **(C-D)** Upset plots showing the overlap between the sets of differentially expressed up-**(C)** and down-regulated **(D)** genes in P60 versus P90 hNPC-treated rods, cones, Müller glia and microglia. **(E-F)** Dot plot showing the selected pathway enrichment results from DEGs that were up- and down-regulated in P60 hNPC-treated rods, cones, Müller glia and microglia compared to P90. **(G-J)** Feature plots showing expression of essential DEGs involved in selected cell-specific pathways in P60 and P90 hNPC-treated rods **(G)**, cones **(H)**, Müller glia **(I)** and microglia **(J)**.

GO analysis of DEGs between P60 and P90 hNPC-treated Müller glia showed upregulation of genes associated with oxidative phosphorylation (*Mt-nd3, Mt-nd5*), synapse assembly, axon guidance (*Sema5a, Sema3a,*) and cAMP-mediated signaling (*Pde4d, Pde10a*) (**Figure 6E, I and S10C**) at P60. Conversely, by P90, they showed increased expression of genes related to phagocytosis (*Cyba, Aif1*), glial cell activation (*Jund, C1qa*), TNF signaling (*Selenos, Zfp36*), synapse pruning (*C1qa, C1qb,*), apoptosis (*Aif1, Ctsd*), metabolic process (*Ckb, Rpsa*) and glucocorticoid response (*Icam1, Zfp36*) (**Figure 6F**). These findings suggest a transition from neuroprotective roles to defensive and reactive states over time with disease progression, aiming to mitigate stress and support retinal homeostasis during degeneration.

At P60, microglia exhibited upregulation of genes related to cell projection assembly (*Nrp1, Slit2, Robo2*) (**Figure 6E**), while by P90, hNPC-treated microglia exhibited increased expression of genes related to lysosome formation (*Tpp1, Ctsd*), antigen processing and presentation (*Ctss, Rt1-Db1*), inflammatory response (*Cebpb, Nlrp3*), T cell activation (*Cd74, Icam1)* leukocyte mediated immunity (*Nlrp3, C3, Il1b)* and superoxide metabolic processes (*Nfe2l2, Ncf1*) (**Figure 6F, J and S10D**). Additionally, the upregulation of genes related to lipid transport (*Abca1, Apoe*) and defense response (*Cebpb, Grn, Il1b*), highlights their role in stress management. These findings suggest that microglia play a supportive role in maintaining retinal neuronal connectivity and function at the early stage of retinal degeneration and are more actively involved in immune surveillance, inflammation, and stress management by removing cellular debris to maintain retinal homeostasis as retinal degeneration progresses.

### Temporal dynamics of cell-cell interactions between grafted hNPCs and retinal cells

To understand how grafted hNPCs interact and communicate with host retinal cells, we investigated the cell-cell interactions and communications over time. We first integrated single-cell datasets from grafted hNPCs with hNPC-treated host retinal cells by mapping one-to-one orthologous genes with high confidence using ENSEMBL multiple species comparison tool^67^ and created UMAP following the SeuratV4 workflow, applying Harmony^68^ for batch-correction (**Figure 7A**). Next, we applied CellChat to compare and identify the signaling pathways mediating the interaction and communication between hNPCs and retinal cells based on the expression of ligand and receptor pairs. This analysis revealed decreased number of signaling interactions (**Figure 7B**) and intercellular communication strength (**Figure 7C**) between hNPCs and host retinal cells at P90 compared to P60 (**Figure 7D-E**). Further, we used an unsupervised machine learning-based approach to analyze the functional changes in the landscape of intercellular communication of specific signaling pathways (**Figure 7F**) and to compute the relative information flow between P60 and P90 retina (**Figure 7G**). We found trophic factors, synaptic, and cell adhesion signaling pathways were downregulated in P90 compared to P60 (**Figure 7H-I and S11-14**). Notably, MANF, FGF, BMP, and JAM signaling were markedly downregulated. MANF and FGF-2 are known to protect photoreceptors.^30,69^ Our analysis revealed increased *Fgf1* and decreased *Manf* and *Fgf2* in the rods of P90 compared to P60 (**Figure 7J, S9D and S15A**). Similarly, the expression of their receptors *Fgfr1, Kdelr1,* and *Kdelr2* was upregulated, and *Nptn* was downregulated in the rods at P90. On the other hand, decreased *Fgf1* and increased *Manf* and *Fgf2* were noted in the cones at P90 (**Figure 7J, S10D and S15B**). We also observed increased *Kdelr1,* decreased *Kdelr2* and *Nptn*, while no change in *Fgfr1* expression in the cones (**Figure 7J, S11D and S15B**). Recent studies have demonstrated that MANF interacts with NPTN to mitigate inflammatory response and apoptosis by inhibiting NF-κB signaling.^38,70^ MANF also serves as an immunomodulatory agent by enhancing tissue repair and regeneration in the retina.^31^ Additionally, MANF has been shown to exert neuroprotective effects to repair brain tissue after ischemic injury by increasing the number of phagocytic innate immune cells.^71^ Interestingly, double immunostaining of IBA1 and MANF confirmed significantly higher expression of *MANF* and increased number of MANF^+^ microglia in the hNPC-treated retina at P60 compared to P90 (**Figure 7K-M**). Further, overall increased expression of *MANF* was detected in P60 hNPC treated retina compared to P90 (**Figure 7K-L**).

**Figure 7:**
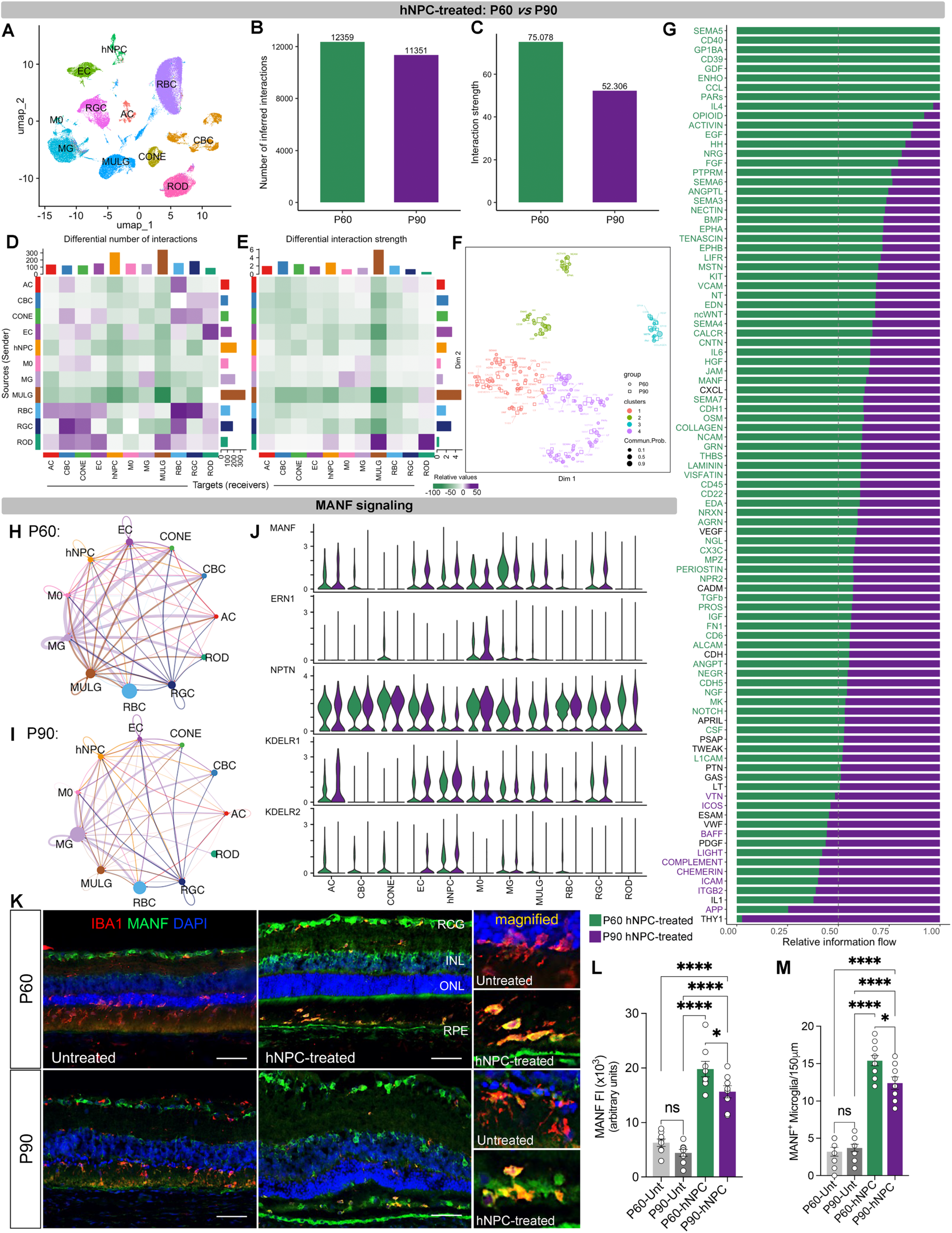
CellChat analysis on cell-cell communication signaling networks identified MANF signaling as a key mediator of hNPC-induced photoreceptor protection. **(A)** UMAP representation of integrated scRNA seq data of *in vivo* hNPCs and rat retinal cells. **(B-C)** Bar plots showing the total number of cell-cell interactions **(B)** and their interaction strengths **(C)** for P60 and P90 hNPC-treated retina. **(D-E)** Heatmap showing the differential number **(D)** and strength **(E)** of cell-cell communication interactions among the host retinal cells and with grafted hNPCs in P60 and P90 hNPC-treated retina. **(F)** Joint dimensionality reduction and clustering of signaling pathways according to their functional similarity inferred from P60 and P90 data. Each dot represents the communication network of one signaling pathway. Dot size is proportional to the overall communication probability. Different colors represent different groups of signaling pathways. **(G)** Bar plots showing signaling pathways with significant differences between P60 and P90 hNPC-treated retina. All significant signaling pathways were ranked based on differences in overall information flow (sum of communication probability among all pairs of cell populations in the network) within the inferred networks between P60 and P90 retina. The top signaling pathways colored green are more enriched in P60 retina, the signaling pathways colored purple are more enriched in P90 retina, and the signaling pathways colored black are non-significant. **(H-I)** Network plots showing the cell-cell communication signaling strength between different cell populations in P60 and P90 hNPC-treated retina for the MANF signaling pathway. Circle sizes proportionate to the number of cells in each cell group, and edge width represents the probability of communication. **(J)** Violin plots showing the expression of *MANF* (ligand) and their receptors (*ERN1, NPTN, KDELR1, and KDELR2*) in P60 and P90 hNPC-treated retinal cells. **(K-L)** Immunofluorescence and quantitative analysis showing significantly increased expression of MANF (green) in P60 and P90 hNPC-treated retinal cells and Iba1^+^ microglia (red) compared to untreated. Scale bar = 50μm. Abbreviations: RPE, retinal pigment epithelium; INL, inner nuclear layer; ONL, outer nuclear layer; RGC, retinal ganglionic cell. Data are represented as mean ± SEM. One-way ANOVA with Tukey’s test was used for multiple comparisons. *\*P ≤ 0.05, ****P ≤ 0.0001, ns: nonsignificant*. **(M)** Bar graph showing increased counts of MANF^+^ microglia in P60 and P90 hNPC-treated retina compared to untreated. Notably, MANF^+^ microglia significantly decreased in P90 hNPC-treated retina compared to P60 hNPC-treated retina. Data are represented as mean ± SEM. One-way ANOVA with Tukey’s test was used for multiple comparisons. *\*P ≤ 0.05, ****P ≤ 0.0001, ns: nonsignificant*.

Further, BMP signaling which induces retinal regeneration and provides neuroprotection by activating Smad signaling and MAPK-mediated upregulation of FGF signaling^72,73^, was significantly downregulated at P90 (**Figure S11G-H**). Our recent study also demonstrated that hNPCs offer vision protection via increasing phosphorylation and activation of AKT-ERK as a key survival pathway.^19^ Cell adhesion molecule (CAM) signaling like NECTIN, NRXN, and NCAM were significantly upregulated at P60 (**Figure S12**), contributing to synaptic stability and visual signal transmission. Notably, NECTIN signaling features in all the retinal cells but not rods, hNPCs, and microglia. By P90, CAM signaling declined, potentially reducing synaptic connectivity between the retinal neurons and disrupting the transmission of visual signals, ultimately impacting visual function.^74–82^ Additionally, COLLAGEN and LAMININ signaling pathways, which are critical for extracellular matrix (ECM) remodeling, structural integrity, and cell-ECM interactions, were significantly downregulated at P90 compared to P60. COLLAGEN signaling supports retinal stability and cellular adhesion, while LAMININ signaling is essential for retinal cell adhesion, differentiation, and survival.^83–85^ The decline in these pathways suggests weakened ECM integrity and diminished support for retinal cell function during disease progression. Despite these declines, new CADM signaling-based communications emerged at P90 between hNPCs and retinal neurons, including rods, cones, and CBCs, which were absent at P60 (**Figure S12A-B**). CADM signaling (CADM1 and CADM3) is essential for visual circuit formation by regulating axon guidance, neurite growth, and synapse formation.^75,86^ Similarly, CDH signaling was reduced at P90 but showed newly established communications between rods and other retinal cells, including cones, RBCs, CBCs, and ACs, partially compensating for declining synaptic pathways (**Figure S14C**). Furthermore, PTN signaling remained stable between hNPCs and retinal cells over time (**Figure S11E-F**). Notably, in addition to hNPCs, microglia, and Müller glia also produced MANF. It is noteworthy that while significantly increased *MANF* and decreased *PTN* were detected in P90 hNPCs compared to P60 hNPCs, overall MANF signaling between hNPCs and retinal cells was downregulated in P90, but PTN signaling was unchanged. Altogether, these findings indicate a progressive decline in critical trophic and growth factors, cell adhesion, and ECM-related signaling pathways, suggesting reduced photoreceptor protection and synaptic stability in the hNPC-treated retinas over time. However, the emergence of new CADM and CDH signaling-based interactions between hNPCs and other retinal cells at P90, partially supports synaptic maintenance and visual function. Together, these results suggest that hNPC-mediated photoreceptor protection evolves over time with distinct shifts in intercellular communication strategies as retinal degeneration progresses.

## DISCUSSION

Stem cell therapy for retinal degenerative diseases has been extensively investigated in preclinical and clinical settings, showing promise for vision preservation. However, efficacy deterioration over time has been reported consistently. The mechanisms underlying this decline as well as the status of grafted stem cells and their interaction with host retinal cells over time, remain poorly understood. Using scRNA-sequencing, we investigated the transcriptomic changes in grafted hNPCs and host retinal cells following subretinal injection in a well-established rodent model for RP. Our findings revealed that grafted hNPCs and host retinal cells undergo dynamic transcriptomic changes as retinal degeneration progresses. Grafted hNPCs contribute to vision protection through multiple modalities, including trophic factor release, metabolic regulation, and reduction of apoptosis, oxidative stress, inflammation, and remodeling of the ECM. However, the gradual decline in the therapeutic efficacy of hNPCs over time might be the cumulative result of decreased trophic factors support from both donor cells and host retina, increased oxidative stress and inflammation, and reduced signaling interactions between grafted hNPCs and host retinal cells within the degenerative environment. The host retinal environment plays a pivotal role in graft survival and function, directly influencing the efficacy of stem cell therapy. These findings provide valuable insights into the dynamic transcriptomic changes in the grafted hNPCs and host retinal cells, emphasizing the need for a comprehensive approach. Combining stem cell transplantation with strategies to improve the host retinal environment—such as supplying trophic factors, mitigating oxidative stress, reducing inflammation, and enhancing ECM integrity—may be essential for achieving long-term vision protection.

We identified that hNPCs express key trophic factors such as MANF, MYDGF, MDK, and PTN, which play an important role in photoreceptor survival and maintenance. These factors mitigate oxidative stress, inflammation, and apoptosis, thereby supporting retinal function and integrity.^31,35,38,70,87–90^ At P60, increased *MDK* and *PTN* in grafted hNPCs likely promote their survival, migration, and differentiation. In contrast, at P90, increased *MANF* and *MYDGF* in hNPCs might serve as a protective response to mitigate increased cellular stress and immune and inflammatory responses due to progressive retinal degeneration, providing additional trophic support to photoreceptors.

Our findings also demonstrate that hNPC transplantation induced distinct transcriptomic changes across retinal cell types. In photoreceptors, hNPCs preserved gene expression patterns associated with visual perception and photoreceptor maintenance, mitigating degeneration and preserving vision. Notably, hNPCs significantly increased the expression of rod-derived cone viability factor (RdCVF: *Nxnl1*) in rods, an essential factor that protects cones from oxidative stress^50^ and promotes their survival by stimulating aerobic glycolysis to support their metabolic demands and maintaining their function.^51^ Degeneration of rods results in the deprivation of RdCVF, leading to cone degeneration in rodent and human RP.^91,92^ By restoring RdCVF expression, hNPC treatment counteracts this degenerative cascade, supporting cone viability and visual function. Preservation of cone function and their connections with second-order neurons following hNPC treatment has been reported in our recent studies.^19^

Inflammation and immune activation are hallmarks of retinal degeneration, and effective immune regulation is critical for stem cell therapy success.^93–95^ Our findings demonstrate that hNPC treatment not only supports photoreceptor survival but also modulates the behavior of retinal supporting cells, such as Müller glia and microglia, contributing to a protective retinal microenvironment. At P60, hNPC treatment promoted microglial surveillance and phagocytosis, aiding in the clearance of shed POS and mitigating oxidative stress—key processes for maintaining retinal homeostasis.^63,96^ By P90, microglia transitioned to phagocytic phenotypes with increased lipid metabolism to mitigate further photoreceptor degeneration.^65,66^ Temporal transcriptomic shifts in Müller glia and microglia over time further suggested compensatory mechanisms aimed at supporting retinal homeostasis. hNPC treatment stimulated Müller glia to upregulate neurotrophic signaling pathways, reinforcing their protective functions. Concurrently, microglia demonstrated reduced expression of pro-inflammatory genes, indicating an hNPC-driven modulation of immune responses to suppress inflammation and prevent photoreceptor apoptosis. Notably, weakened Müller-microglial interactions in untreated retinas were associated with increased photoreceptor apoptosis and impaired retinal homeostasis^41^. In contrast, hNPC treatment appeared to restore the bi-directional crosstalk between Müller glia and microglia, enabling a coordinated response to support retinal integrity and slow the progression of retinal degeneration.^97–99^ Together, these findings highlighted the ability of hNPCs to influence retinal supporting cells, fostering a homeostatic environment conducive to photoreceptor survival.

Using CellChat, we analyzed hNPC-induced changes in cellular communication signaling among retinal cells over time, identifying trophic factor (MANF, FGF, BMP), synaptic (NRXN, NCAM), cell adhesion (CADM, CDH), and ECM-related (COLLAGEN, LAMININ) signaling pathways as primary mechanisms of photoreceptor protection. All these cell-cell communication pathways were highly upregulated at P60, providing trophic support, synaptic stability, and ECM integrity, essential for photoreceptor survival and maintaining retinal homeostasis. However, by P90, the downregulation of these pathways likely reflects a progressive decline in trophic factor support, synaptic stability, and ECM integrity, resulting in reduced photoreceptor protection. Additionally, downregulated CAM signaling potentially impairs synaptic connectivity and visual signal transmission at P90. Despite these declines, adaptive mechanisms emerged at P90. hNPCs established new signaling interactions with retinal cells, particularly rods and cones, *via* CADM and CDH signaling, to partially compensate for the photoreceptor protection. CADM signaling facilitated synaptic organization and visual circuit stability, while strengthened CDH signaling between rods, hNPCs, and Müller glia supported photoreceptor survival over time. However, the overall reduction in cell-cell signaling interaction strength and intercellular connections at P90 likely contribute to the progressive decline in hNPC therapeutic efficacy as retinal degeneration advances. These findings highlight the dynamic changes in hNPC-induced cellular communication, emphasizing the need for strategies to sustain trophic support and ECM-remodeling to preserve retinal integrity and maximize long-term therapeutic outcomes.

Our study comprehensively explores the temporal gene expression dynamics in grafted hNPCs and their therapeutic impact on host retinal cells within the degenerative retinal environment. Grafted hNPCs survived, migrated, and expressed multiple trophic factors, effectively stimulating host retinal rods, cones, Müller glia, and microglia to promote neuroprotection and retinal homeostasis. Overall, grafted hNPCs dynamically adapt to the degenerative retinal environment, exhibiting shifts in differentiation, metabolic activity, and immune response over time. Concurrently, hNPC transplantation modulates host retinal cell behavior, enhancing neuroprotection and reducing inflammatory responses. These findings underscore the therapeutic potential of hNPCs in mitigating retinal degeneration and preserving photoreceptor function, highlighting them as a promising strategy for the treatment of retinal degenerative diseases.

## Limitations of the study

While this study provides valuable insights into the transcriptomic remodeling of grafted hNPCs and host retinal cells over time (P60 and P90), additional later time points are needed to capture the temporal dynamics of these changes. A more comprehensive time-course analysis of host retinal degeneration will provide details of the molecular changes of photoreceptors and retinal supporting cells, which will assist in identifying the best time of windows for intervention. Another limitation is a single subretinal injection of stem cells only affects about one-third of the retinal area, leaving the rest of the retina undergoing progressive retinal degeneration. Multiple injections covering the majority of the retinal area will reduce the negative impact of the degenerative retinal environment and promote donor cell survival and function.

## ACKNOWLEDGMENTS

California Institute Regenerative Medicine (LSP1-08235). J.C. was supported by CIRM-EDUC-08383 and S.R. was supported by CIRM-EDUC2-12638, and funding from the Board of Governors Regenerative Medicine Institute at Cedars-Sinai Medical Center.

## AUTHOR CONTRIBUTIONS

S.S., S.W. conceptualized this study; S.S., B.L. S.W., H.X., J.S.A., S.R., and J.C. performed animal surgery experiments and collected samples; S.S., B.L, and S.W. performed single-cell isolation of hNPCs and rat retinal cells. B.L. and J.C. performed ERG and OKR; S.S., S.R., and J.C. performed cryosectioning and immunostaining; S.S. and S.BA. performed bioinformatics analysis of hNPC datasets; S.S., S.BE., and S.BA. performed bioinformatics analysis of rat datasets; S.S. prepared the Figures with assistance from S.BA.; S.S. drafted the manuscript; S.S., S.W., B.L., S.SV, C.N.S., S.BA., A.L., and V.S. edited the manuscript. All authors have read and approved the manuscript.

## DECLARATION OF INTERESTS

The authors declare no conflict of interest.

## METHODS

### EXPERIMENTAL MODEL AND SUBJECT DETAILS

#### Ethical statement

All animal experimental procedures were approved by Cedars-Sinai Medical Center’s Institutional Animal Care and Use Committee (IACUC 7611) and the ARVO Statement for the Use of Animals in Ophthalmic and Vision Research.

#### Animal model

RCS rats were maintained in a 12 h light/dark cycle at 20–25 °C and controlled humidity (30– 70%) in a pathogen-free facility with ad libitum access to food and water. Both male and female rats were used for all experiments. The rats between postnatal day (P) 21-23 were randomly assigned to two groups: one group received a subretinal injection of human neural progenitors (hNPCs) (n=65), while the other served as untreated controls (n=44). To minimize the immune response to human-derived cells in this xenograft model, cyclosporine A was administered via drinking water (210 mg/L), starting 1 day prior to transplantation and continuing until sacrifice. The experiment was repeated 3 times for P60 (untreated: n=24; hNPC-treated: n=30) and 2 times for P90 (untreated: n=20; hNPC-treated: n=35) timepoint.

### METHOD DETAILS

#### Cell Preparation and Transplantation

The generation and expansion of hNPCs have been previously described.^100,101^ Research-grade hNPC was used in this study under the Stem Cell Research Oversight Committee (Pro00025772) at Cedars-Sinai Medical Center.

Cells were prepared according to our published protocol.^102^ Briefly, hNPC vials were rapidly thawed in a 37 °C water bath after removal from liquid nitrogen, followed by resuspension in prewarmed Stemline medium (Sigma, St. Louis, Missouri) supplemented with rhEGF (100 ng/mL; Millipore, Billerica, MA) and rhLIF (100 ng/mL; Millipore). The cell suspension was centrifuged at 200g for 5 minutes, and the pellet was resuspended in Balanced salt solution (BSS, Alcon). Cells were then labeled using PKH26 Red Fluorescent Cell Linker Kits (Sigma, St. Louis, Missouri) according to the manufacturer’s protocol, and finally resuspended at a concentration of 30,000 cells/μL in BSS. Cells were kept on ice until transplantation.

Cell injections were performed following a previously established protocol.^102^ Briefly, 2 μL of the hNPC suspension (6 × 10^4 cells/eye) was injected into the subretinal space of both left and right eyes through a small scleral incision using a fine glass pipette (internal diameter 75-150μm) attached via tubing to a 25-μL syringe (Hamilton). To reduce intraocular pressure and minimize cell efflux, the cornea was punctured prior to injection. Immediately after injection, the fundus was examined for retinal damage or signs of vascular distress. Animals showing such issues were excluded from further study.

#### Efficacy Evaluation

##### Optokinetic Response (OKR)

OKR testing provides a non-invasive method for assessing spatial visual acuity, measured in cycles per degree (c/d). Visual acuity was evaluated at P60 and P90, following our previously published protocol.^14,102^ The OKR results were observed and recorded by two independent, blinded investigators. Data were analyzed after incorporating our previous study datasets.

##### Electroretinography (ERG)

Electroretinography (ERG) measures the overall electrical activity of the retina in response to light stimulation. ERG recordings were performed at P60 and P90 according to our previously established protocol.^39,102^ The eyes were stimulated with full-field light flashes using a computer-controlled Espion system (Diagnosys LLC). A total of 20-30 sweeps were recorded for each animal, and the averaged responses were used to determine the response amplitude. Data were analyzed after incorporating our previous study datasets.

#### Single-cell RNA sequencing

Rats were sacrificed at P60 and P90 time points after visual function tests. Eyes were enucleated, and retinas were dissected in AMES solution (equilibrated with 95% O2/5% CO2). Under fluorescent dissecting microscope, red fluorescent region containing injection site (dorsal-temporal part) was dissected from the retina. For untreated retinas, the dorsal-temporal part corresponding to the same location in the cell-treated eye was dissected. Retinas from 8-15 rats were pooled, digested with papain, and dissociated into single-cell suspensions through manual trituration in ovomucoid solution.^103^ Single-cell suspensions from hNPC-treated and untreated retinas were labeled with DAPI. PKH26-labeled hNPCs, along with hNPC-treated and untreated retinal cells, were sorted using a FACS sorter (FACSAria III, BD Biosciences, Franklin Lakes, NJ) and collected into 1.7 mL RNase-free tubes (Eppendorf, Hamburg, Germany) containing 2% BSA in PBS. From hNPC-treated cell suspension, PKH26-labeled hNPCs and treated retinal cells were collected in separate tubes. The sorted cell suspensions were immediately processed for single-cell RNA sequencing following the 10x Chromium v3 single-cell reagent kit protocol for library preparation. cDNA library quantification and quality were assessed with D1000 or D5000 HS kit for the Agilent TapeStation 4200. Libraries were sequenced using Illumina Novaseq 6000 S4 platform using 100bp paired-end sequencing for a sequencing depth of 50,000 read pairs/cell.

#### Histology

Eyes were enucleated from P60 and P90 time points, fixed in 4% paraformaldehyde, and embedded in OCT for cryostat sections. Retinal cryostat sections were placed on a series of 5 slides, with 4 sections per slide. The first slide of each series was used for cresyl violet staining to visualize retinal lamination and donor cell location. The remaining slides were then used for immunofluorescent staining following previously described protocols.^39^ Images were taken with a Leica microscope.

#### Immunofluorescence Staining

Retinal sections were stained with the primary antibodies (KEY RESOURCES TABLE) with our published protocols^39,102^. Anti-mouse or rabbit secondary antibodies conjugated to Alexa Fluor-488 or Alexa Fluor-568 were used and counterstained with 49,69-diamidino-2-phenylindole (DAPI) before mounting slides using Fluorescent Mount media (Sigma-Aldrich).

#### Image Analysis

Differences in immunoreactivity of MANF between untreated and hNPC-treated retina at P60 and P90 were assessed as fluorescent intensities (FI) with our previous protocol.^103,104^ In brief, for FI measurements, MANF positive region of interest (ROI) was selected using ImageJ freehand tool. Fluorescent intensities of ROIs were measured after normalizing to the background. Fluorescent intensity values were averaged (4 sections per retina and 8 retinas per group; each section 40μm apart) to determine signal density. The FI values were presented as arbitrary units. The processing and analysis were kept consistent between images. Statistical analysis was performed on average FI values (mean ± SEM). MANF^+^ microglia were counted manually (4 sections per retina and 10 retinas per group; each section 40μm apart). Image analysis was performed blindly and separately by 2 observers.

#### Preliminary processing of hNPCs and rat retinal cells scRNA-seq raw data

Cell Ranger software (version 4.0.0) (https://support.10xgenomics.com/single-cell-gene-expression/software/pipelines/4.0/what-is-cell-ranger) was used to perform preliminary processing of single-cell sequencing data, including aligning sequencing reads to the reference transcriptome (hNPCs: GRCh38; rat: Rnor6), estimating and filtering, and generating feature-barcode matrices. Wildtype rat sequencing data from the Wang et al., 2022 were downloaded from GEO database (GSE209872: GSM6403183 and GSM6403184).

#### Removing low-quality cells and identification of cell types

Seurat (version 4.4.0) software was used to perform downstream quality control, integration, identification of cell types, and subsequent differential expression gene analysis on the gene-cell expression matrices obtained above.^105^ For hNPCs dataset, cells with less than 250 and greater than 20,000 UMIs, less than 200 genes, and mitochondrial gene percent greater than 20% were excluded as low-quality cells. For rat retinal cell datasets, cells with counts, genes, mitochondrial gene percentage, and ribosomal gene percentage outside of three standard deviations from the mean were excluded as low-quality cells.

For hNPCs, the ‘LogNormalize’ function was used for normalization and scaling of the expression matrix for each sample, and then the ‘SelectIntegrationFeatures’ and ‘FindIntegrationAnchors’ functions were used to select the features and anchors for downstream integration. These anchor points were then used to integrate all datasets of the hNPCs through the ‘IntegrateData’ function. Next, the ‘ScaleData’ and ‘RunPCA’ functions were used for scaling and calculating principal component analysis (PCA) dimensions in the hNPCs integrated datasets. Unsupervised clustering was then performed using the ‘FindNeighbors’ and ‘FindClusters’ functions, grouping the datasets into distinct clusters. Dimensionality reduction was conducted using the ‘RunUMAP’ function. Clusters were annotated based on known retinal cell type markers and hNPC-specific gene markers. Subsequently, hNPC clusters were subset and reprocessed following the same steps described above. The ‘FindAllMarkers’ and ‘FindConservedMarkers’ functions were used to identify and annotate specific markers for the hNPC clusters.

Similarly, for retinal cells, the expression matrix for each sample was normalized and scaled using the ‘SCTransform’ function^106,107^. This was followed by feature and anchor selection with the ‘PrepSCTIntegration’ and ‘FindIntegrationAnchors’ functions. The identified anchors were then used to integrate all datasets from the same tissue using the ‘IntegrateData’ function. Subsequently, the ‘ScaleData’ and ‘RunPCA’ functions were employed to scale the data and perform principal component analysis (PCA) on the integrated datasets of hNPCs and rat retinal cells. Unsupervised clustering of the datasets into distinct clusters was performed using the ‘FindNeighbors’ and ‘FindClusters’ functions. Dimensionality reduction was carried out using the ‘RunUMAP’ function. To investigate the gene expression profile of each cluster, the ‘FindAllMarkers’ function (with avg_log2FC > 0.4) was used to identify highly expressed genes (marker genes) that are specific to each cluster and correspond to distinct cell types based on marker gene expression.

#### Differential gene expression and functional analysis

To identify differentially expressed genes (DEGs) between *in vitro* and *in vivo* hNPCs at P60 and P90, the ‘FindMarkers’ function with the MAST^108^ test in Seurat was applied. DEGs between wildtype, untreated, and hNPC-treated retinal cells, across different retinal cell types were identified using the ‘FindMarkers’ function with the Wilcoxon test. For hNPCs, we used the EnrichR^109^ R package (version 3.2) and the ‘gseGO’ and ‘gseKEGG’ functions from the clusterProfiler^110^ R package (version 4.10.1) to identify enriched biological pathways and processes in our sets of DEGs, as well as in DEGs shared across different differential expression analyses. For rat retinal cells, we employed the ‘gseGO’ and ‘gseKEGG’ functions from the clusterProfiler R package to determine enriched biological pathways and processes in our DEG sets.

#### Cell-cell ligand-receptor interaction analysis

We performed cell-cell ligand-receptor interaction in our scRNA-seq integrated rat retinal cell and hNPC datasets with CellChat^111^ (v 1.6.1), using cell type labels. The human CellChatDB ligand-receptor interaction database was used for this analysis. To facilitate downstream comparisons of the signaling networks in untreated versus hNPC-treated and P60 versus P90 hNPC-treated groups, we ran the CellChat workflow separately on each dataset from different time points. The CellChat object was created using the normalized gene expression matrix and the cell type annotations, removing any cell groups with fewer than 10 cells. We then ran the recommended CellChat workflow using the following functions: identifyOverExpressedGenes, identifyOverExpressedInteractions, projectData, computeCommunProb, filterCommunication, subsetCommunication, computeCommunProbPathway, aggregateNet, and netAnalysis_computeCentrality. We separately merged P60 and P90 untreated and hNPC treated and P60 and P90 hNPC-treated CellChat objects into one object using the mergeCellChat function. We compared the signaling networks separately at P60 and P90 between untreated and hNPC-treated groups and then between P60 and P90 hNPC-treated groups both functionally and structurally using the computeNetSimilarityPairwise function. Further, we used the rankNet function to compute the relative information flow changes between the groups across all signaling pathways. We identified differentially expressed ligands and receptors as well as their signaling pathways using the identifyOverExpressedGenes function, visualizing results with the netAnalysis_signalingRole_heatmap.

### QUANTIFICATION AND STATISTICAL ANALYSIS

All data were analyzed using GraphPad Software’s PRISM (version 10), and the results are presented as the mean ± SEM. Results were analyzed by One-way ANOVA coupled with a Tukey’s multiple comparisons test for more than 2-group. For ERG and OKR, One-way ANOVA, Kruskal-Wallis test, coupled with Dunn’s Multiple comparison test were used. *P* value < .05 was considered significant.

## Notes

### Competing Interest Statement

The authors have declared no competing interest.

### Summary of Updates

Typos were corrected in the manuscript file.

