## Supplemental Figures for "Dynamic Transcriptomic Remodeling in Human Neural Progenitor Cells Reveals Mechanisms for Vision Preservation in Retinitis Pigmentosa Model"

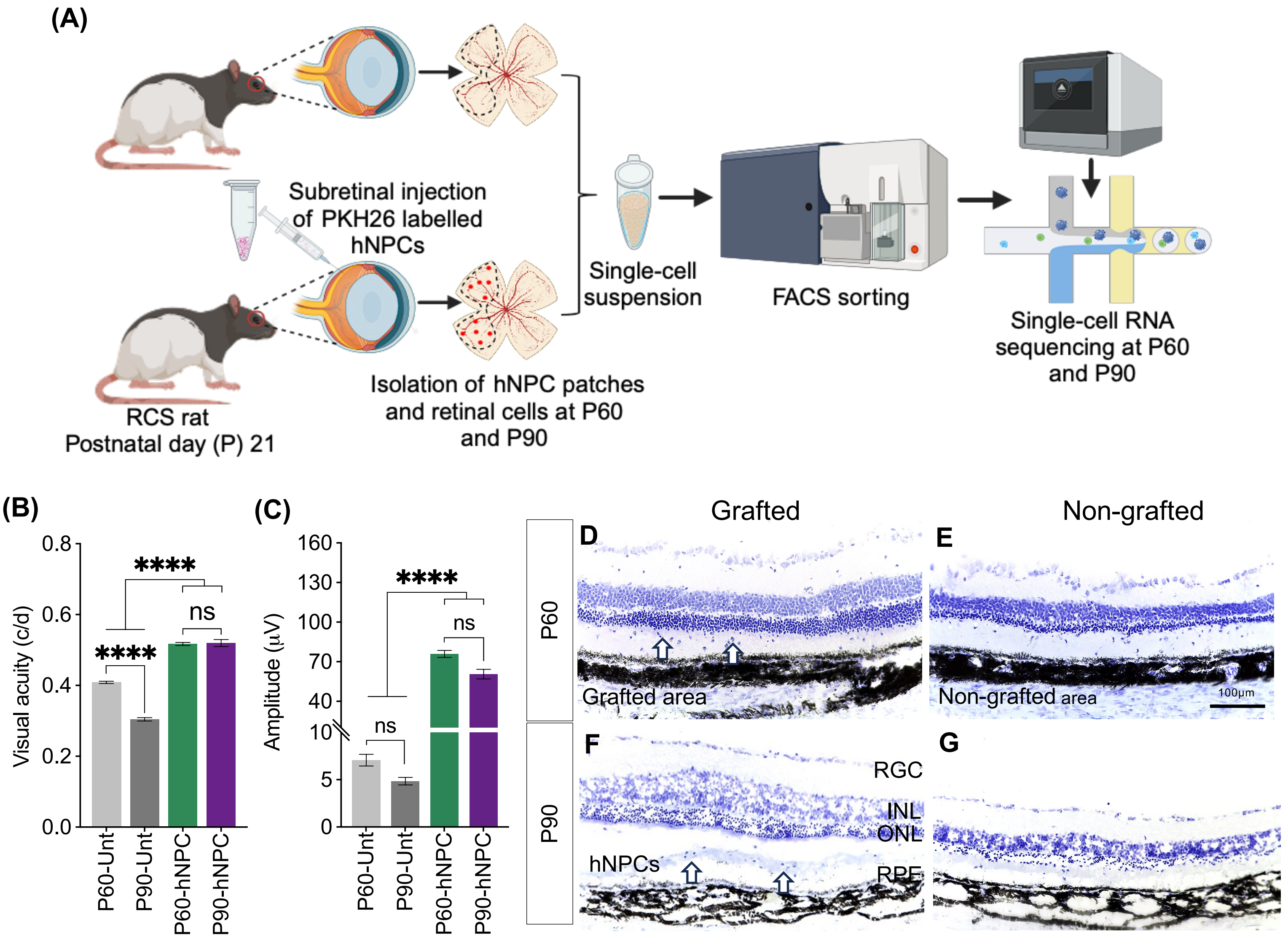


**Figure S1. Experimental protocol, visual function assessment and histological. (A)** Experimental protocol. **(B)** Bar graph of visual acuity tested by Optokinetic response (OKR) on hNPC-treated and untreated RCS rats over time showed significant difference between hNPC-treated and untreated controls at both time points. Notably, visual acuity remains unchanged from P60 to P90 with hNPC-treated eyes, while untreated eyes also showed significant difference between P60 *vs* P90 time points (P60 Untreated: n=217, P60 hNPC-treated: n=121, P90 Untreated: n=48, P90 hNPC-treated: n=66). **(C)** Bar graph of electroretinography (ERG) from P60 and P90 untreated and hNPC-treated RCS rats retina showed significantly higher b-wave amplitude for hNPC-treated eyes compared to untreated controls, while there was no difference between P60 and P90 untreated controls (P60 Untreated: n=105, P60 hNPC-treated: n=120, P90 Untreated: n=85, P90 hNPC-treated n=49). Data are represented as mean ± SEM. One-way ANOVA, Kruskal-Wallis test with post-hoc Dunn's test was used for multiple comparisons. *****P ≤ 0.0001, ns: nonsignificant.*

**(D-G)** Histological evaluation of P60 and P90 hNPC-treated RCS rat retina showed hNPC-treated retinas have thicker photoreceptor layers (P60: 8-12 layers; P90: 6-8 layers) close to the cell-injected side (Arrows in **D&F**) compared to the area away from the graft where 3-4 layers of photoreceptors at P60 **(E)** and 1-2 layers at P90 **(G)** were documented. Scale bar=100µm. Abbreviations: hNPCs: Human Neural Progenitor cells; INL: Inner nuclear layer; ONL: Outer nuclear layer; RGC: Retinal ganglion cell; RPE: Retinal pigmental epithelium.

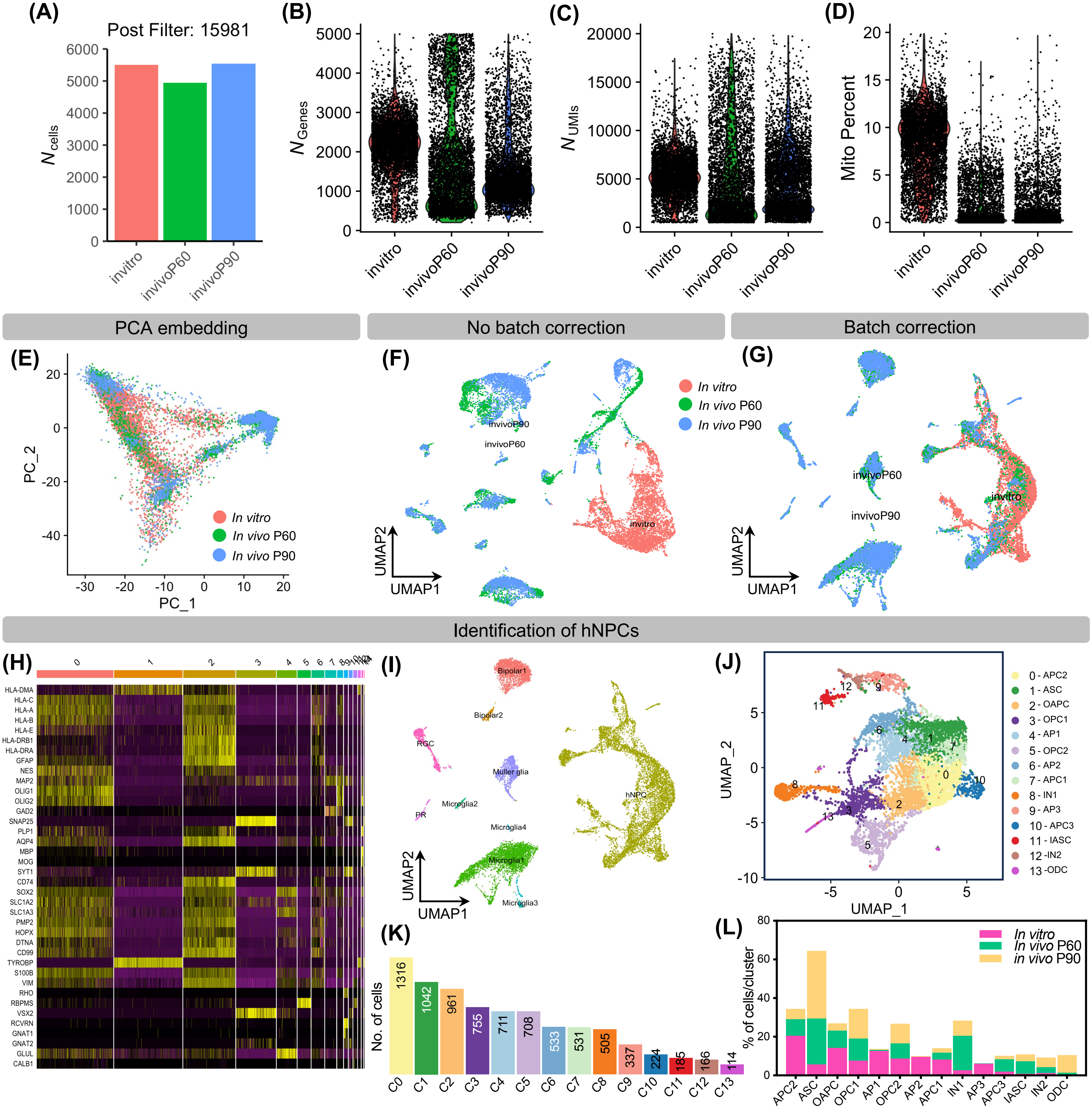


**Figure S2. Quality control of the hNPC scRNA transcriptomics datasets. (A)** Bar plot showing the number of cells in *in vitro* and grafted P60 and P90 hNPCs after quality control. (**B-D)** Violin plots showing the total number of genes, UMIs, and the percentage of mitochondrial genes detected after quality control. (**E**) Scatter plot of PCA embedding. **(F-G)** scRNA-seq UMAPs before (left) and after CCA batch correction (right), colored by experimental batch. **(H)** Heatmap showing the cell-specific marker genes for each cluster. **(I)** UMAP visualization of cell types in the integrated dataset. **(J)** UMAP visualization of hNPC-subpopulations. **(K)** Bar plot showing the number of cells in each subpopulation of hNPCs. **(L)** Bar plot showing the percentage of cells in each subpopulation of *in vitro* and grafted P60 and P90 hNPCs.

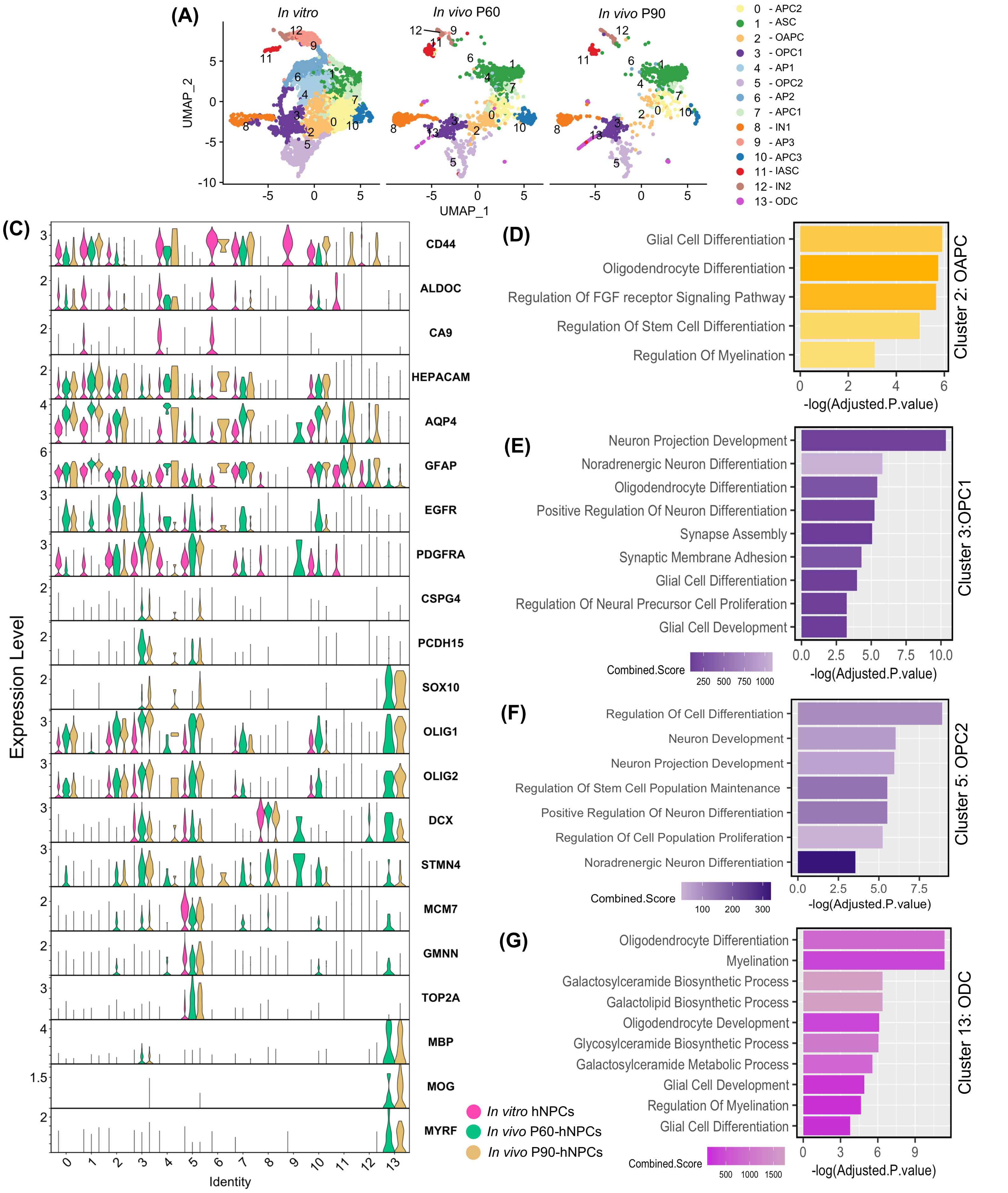


**Figure S3. Characterization of hNPCs. (A)** Bar plot showing the number of *in vitro* and *in vivo* P60 and P90 hNPCs. **(B)** UMAP visualization of *in vitro* and *in vivo* P60 and P90 hNPC-subpopulation clusters colored by cell types. (**C**) Violin plots of cell-type specific marker genes. (**D-G**) Bar plots showing the enrichment of selected upregulated GO-terms related to oligodendrocyte-astrocyte progenitor cells (OAPC; cluster: 2), oligodendrocyte-precursor cells (OPC1; cluster: 3 and OPC2; cluster: 5) and oligodendrocytes (ODC; cluster: 13).

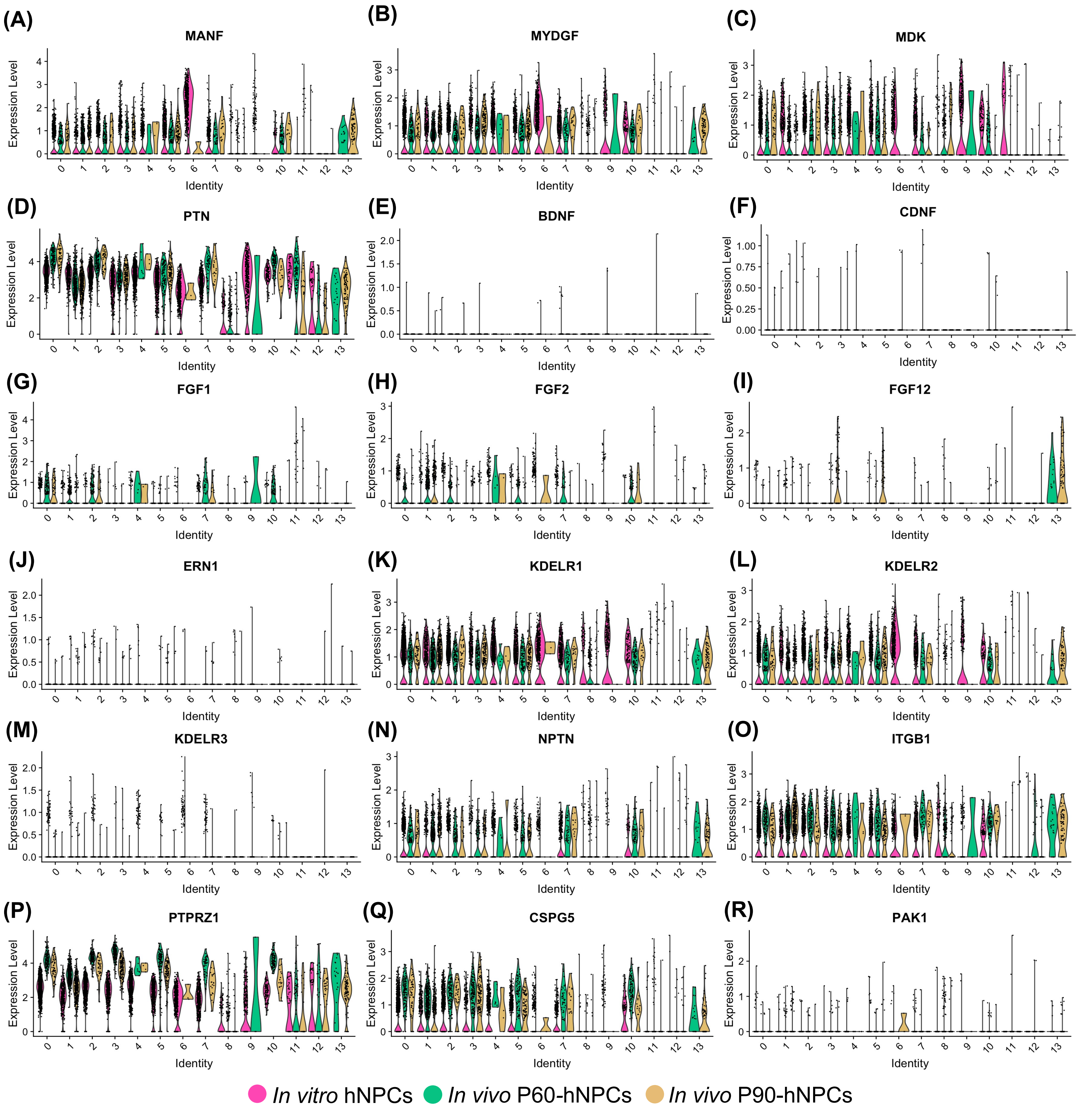


**Figure S4. Expression of trophic and growth factors and their receptors in hNPC-subpopulations. (A-I)** Violin plots showing the expression of trophic and growth factors in all the subpopulations of *in vitro* and *in vivo* P60 and P90 hNPCs. **(J-R)** Violin plots showing the expression of trophic and growth factor receptors in all the subpopulations of *in vitro* and *in vivo* P60 and P90 hNPCs.

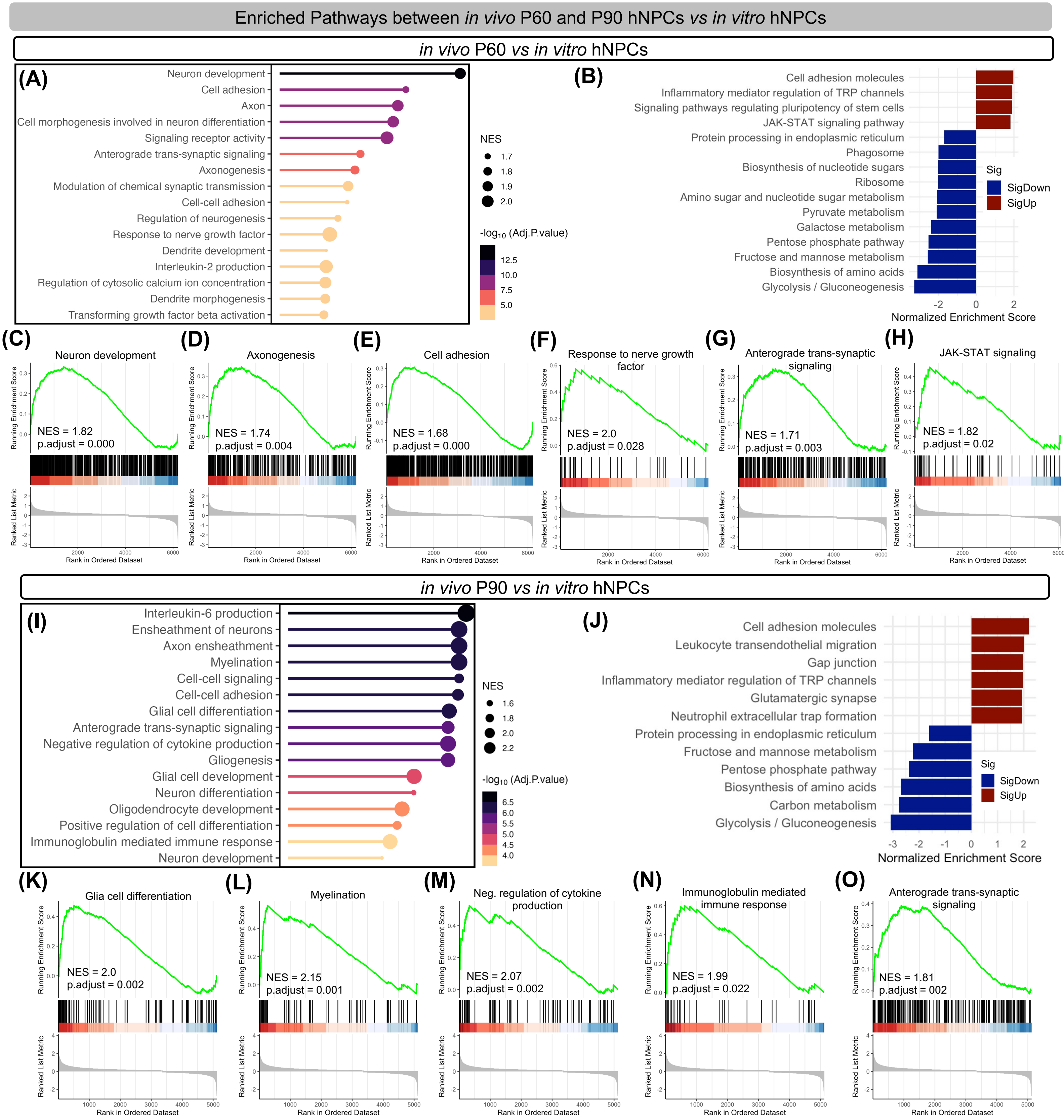


**Figure S5. Transcriptional profiling of *in vivo* P60 and P90 hNPCs compared to *in vitro* hNPCs.**
**(A)** Lolipop plot showing GO biological processes enriched in genes upregulated in P60-hNPCs compared to *in vitro* hNPCs. (**B**) Bar graph showing KEGG pathways enriched in genes up- and downregulated in P60-hNPCs compared to *in vitro* hNPCs. **(C-H)** GSEA plots demonstrating the enrichment of gene sets related to significantly upregulated pathways of P60-hNPCs compared to *in vitro* hNPCs. **(I)** Lolipop plot showing GO biological processes enriched in genes upregulated in P90-hNPCs compared to *in vitro* hNPCs. (**J**) Bar graph showing KEGG pathways enriched in genes up- and downregulated in P90-hNPCs compared to *in vitro* hNPCs. **(K-O)** GSEA plots demonstrating the enrichment of gene sets related to significantly upregulated pathways of P90-hNPCs compared to *in vitro* hNPCs. NES, normalized enrichment score.

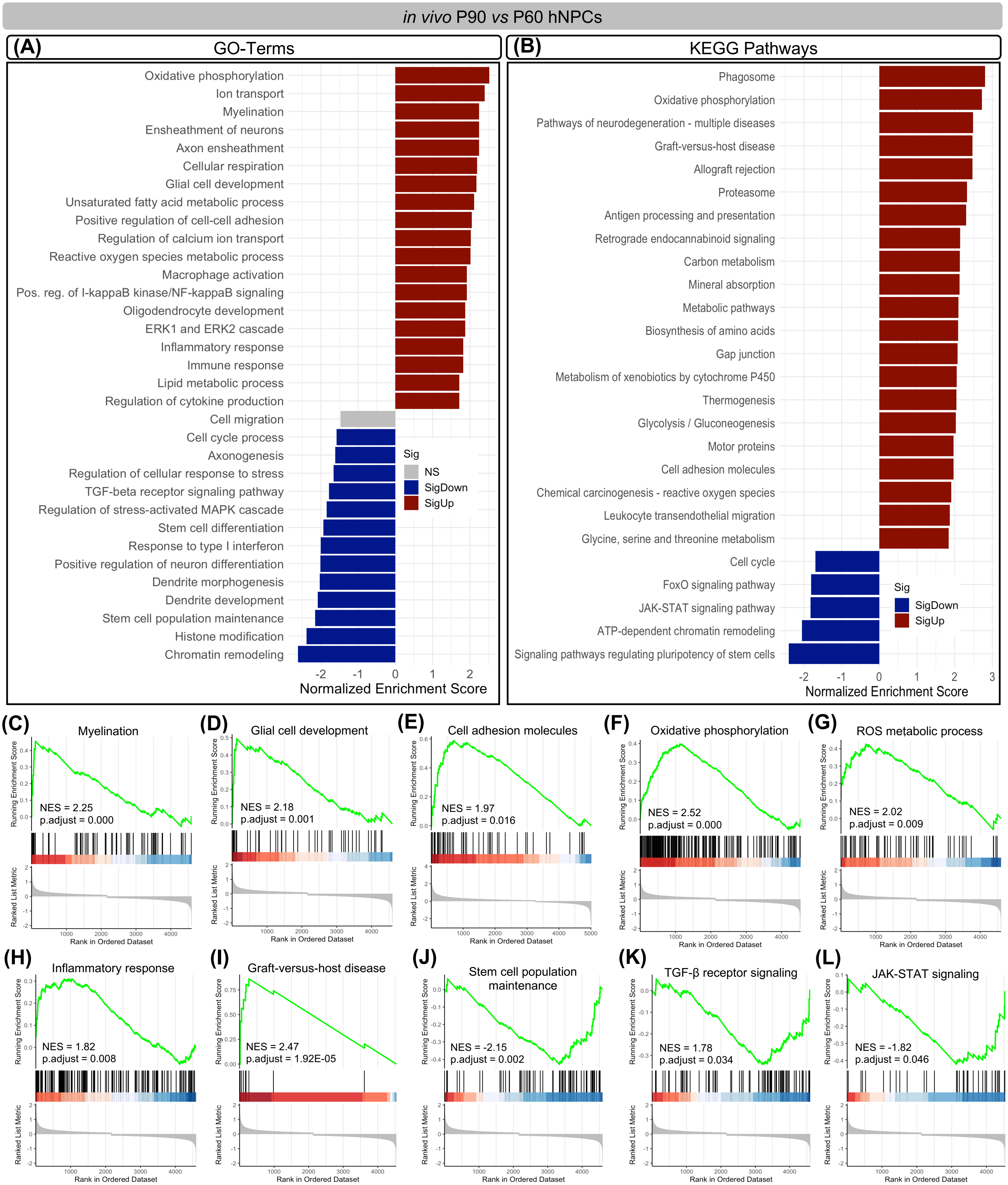


**Figure S6. Transcriptional profiling of *in vivo* P90-hNPCs compared to P60-hNPCs.**
**(A-B)** Bar graphs showing GO biological processes **(A)** and KEGG pathways **(B)** enriched in genes up- and downregulated in grafted P90-hNPCs compared to P60-hNPCs. **(C-L)** GSEA plots demonstrating the enrichment of gene sets related to significantly upregulated **(C-I)** and downregulated **(J-K)** pathways of P90-hNPCs compared to P60-hNPCs. NES, normalized enrichment score.

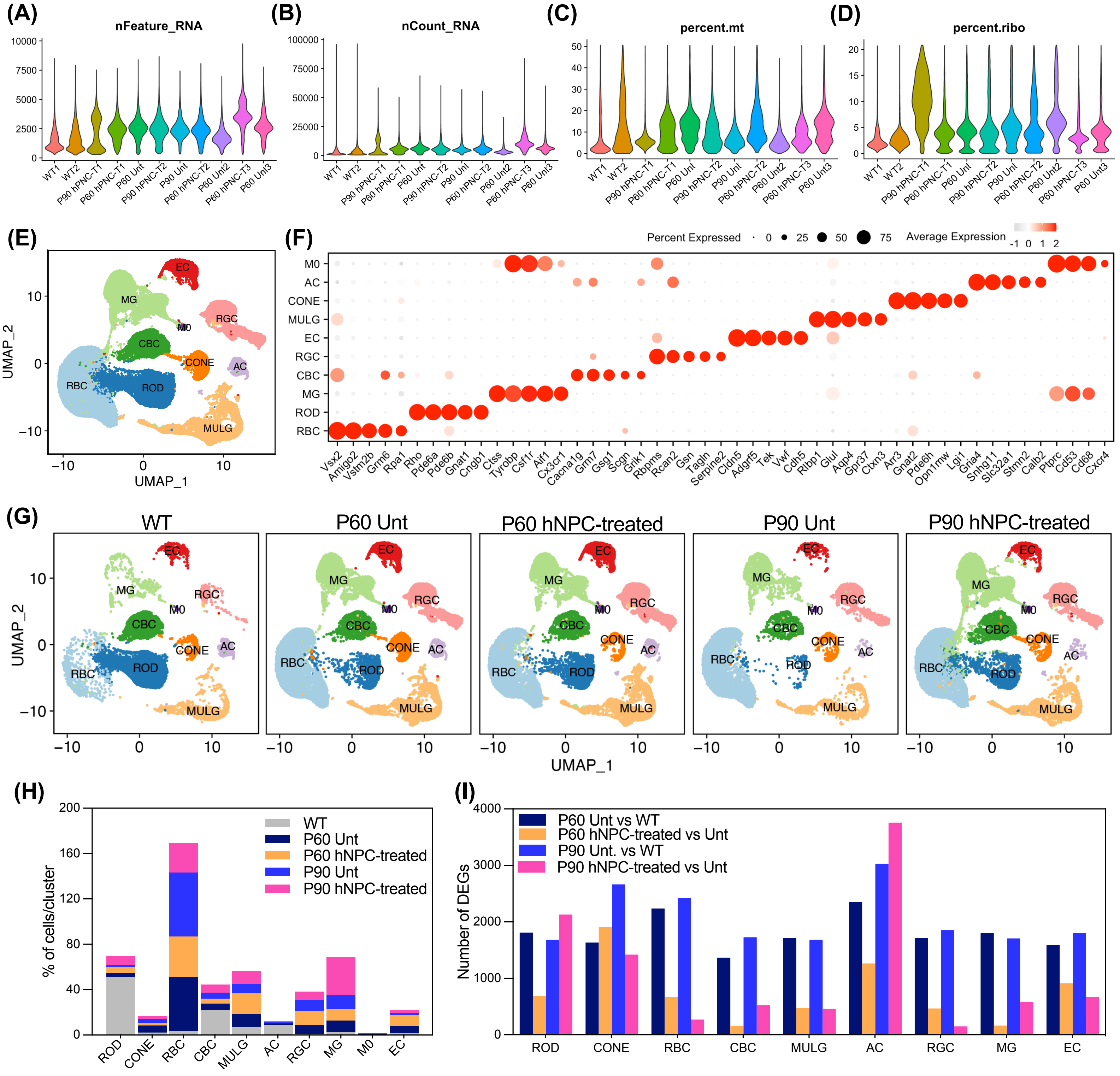


**Figure S7. Quality control of rat retinal scRNA transcriptomics. (A-B)** Violin plots showing distributions of features and counts in rat retinal samples. **(C-D)** Violin plots showing the percentage of mitochondrial and ribosomal genes in rat retinal samples. (**E**) UMAP plot of 118,478 rat retinal cells from wildtype (WT), untreated (P60 and P90), and hNPC-treated RCS (P60 and P90) retina showing partition-based Leiden clustering of different retinal cell types, colored by their cell type-specific annotated identities. **(F)** Dot plot showing the average gene expression of cell type-specific marker genes for the transcriptomically defined clusters. **(G)** UMAP visualization of all the retinal cell types from each group. **(H)** Bar plot showing the percentage of different retinal cell types in each group. **(I)** Bar plot showing the number of differentially expressed genes (DEGs) for each cell type among different groups.

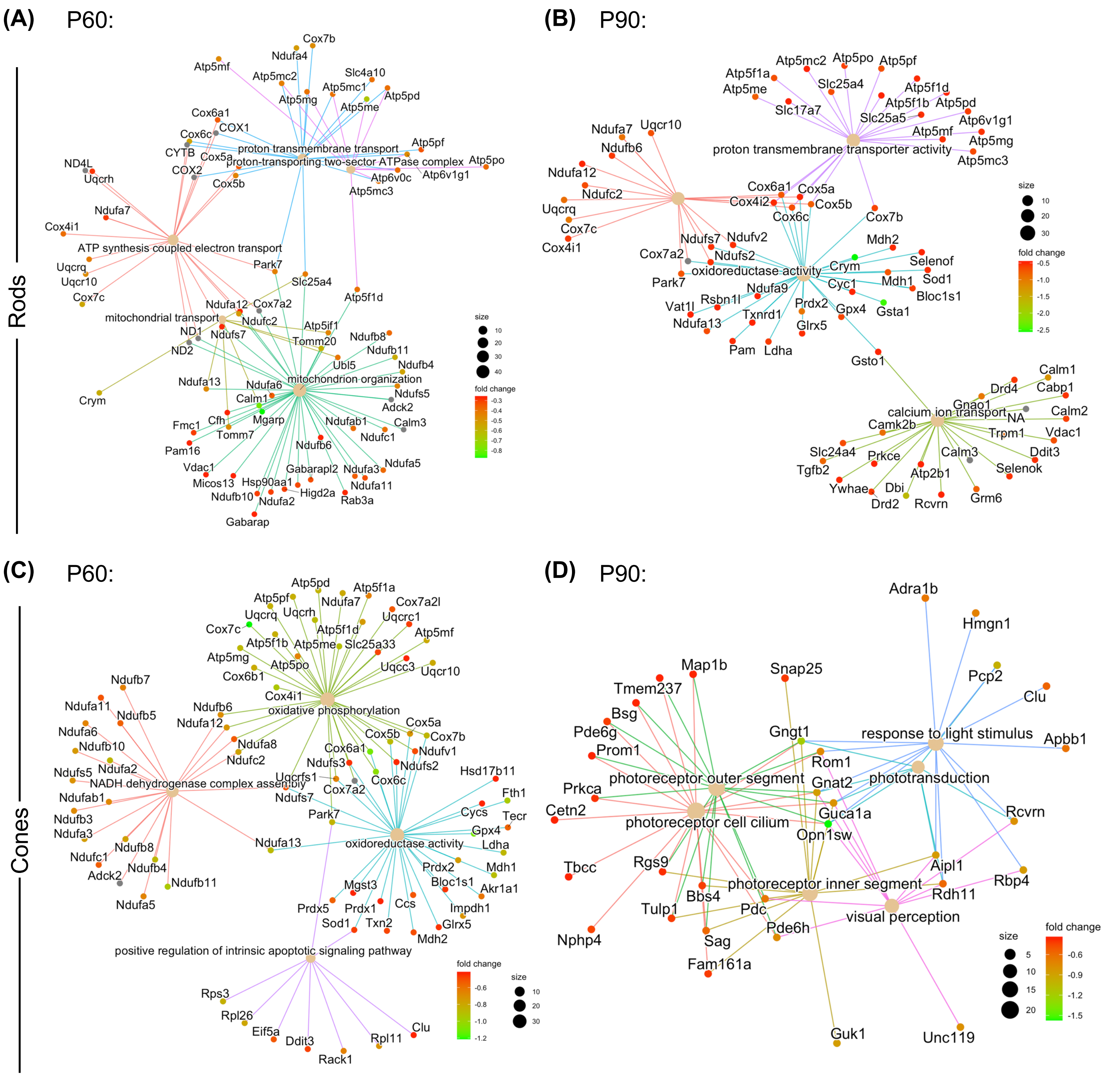


**Figure S8. Downregulated pathways in hNPC-treated rods and cones. (A-D)** Cnet plots showing the interactions of selected downregulated pathways, and their associated genes in P60 and P90 hNPC-treated rods **(A and B)** and cones **(C and D)** compared to untreated. The size of each circle associated with each pathway represents the number of genes enriched in each pathway, and the color of each gene represents the $\log_2$fold change (fold change).

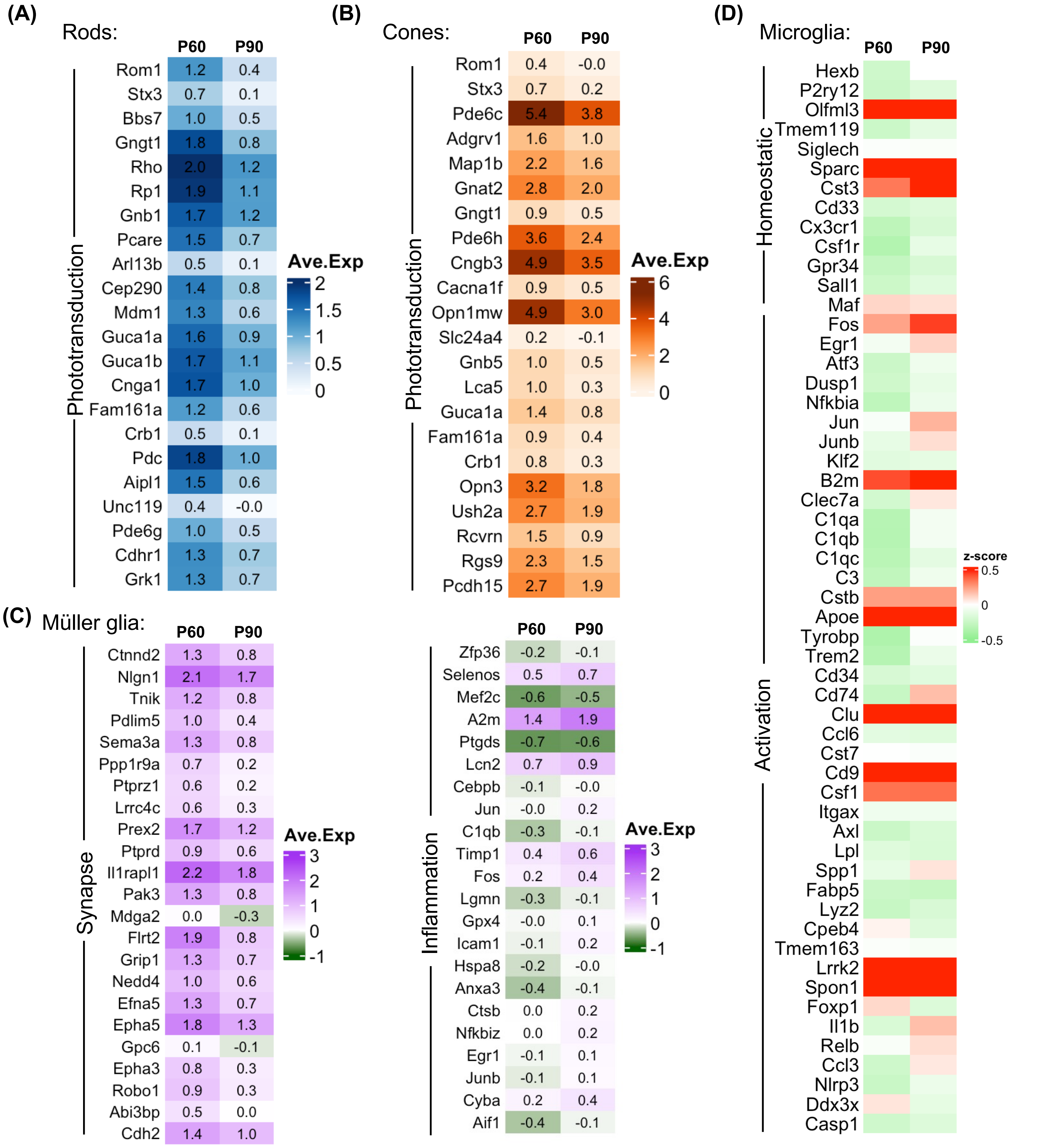


**Figure S9. Selected pathway-specific genes in rods, cones, Müller glia and microglia. (A-D)** Heatmaps showing the expression of selected phototransduction genes in P60 and P90 hNPC-treated rods **(A)** and cones **(B),** synapse formation and inflammatory genes in Müller glia **(C),** and homeostatic and activation (early and late disease-associated) genes in microglia **(D).**

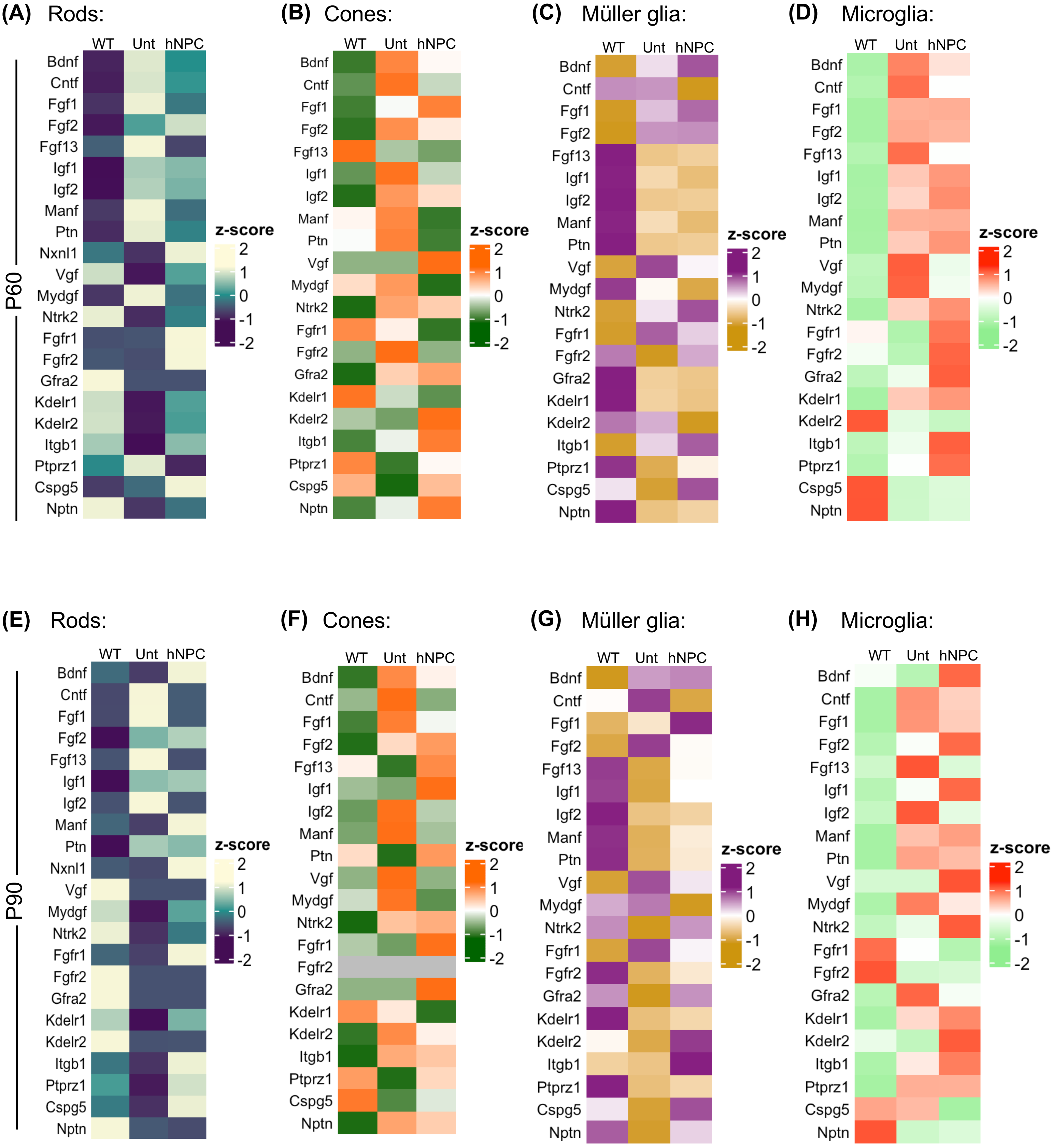


**Figure S10. Expression of neurotrophic and growth factors as well as their receptors in wild-type, untreated, and hNPC-treated RCS rat retinal cells at P60 and P90. (A-H)** Heatmaps showing the expression of different neurotrophic and growth factors as well as their receptors in rods **(A and E)** cones **(B and F)**, Müller glia **(c and G)**, and microglia **(D and H)**, in wild-type, untreated, and hNPC-treated retina at P60 and P90.

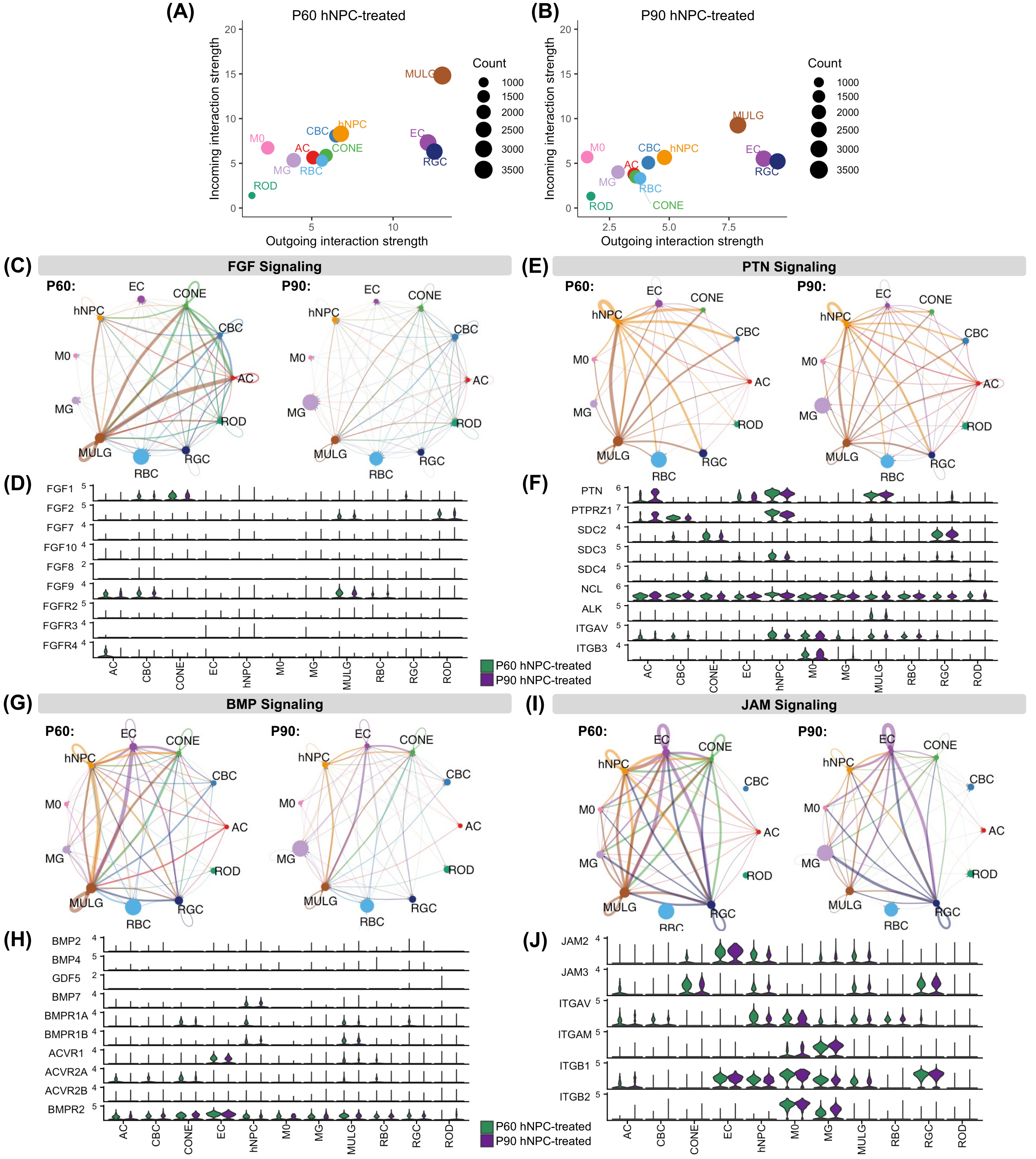


**Figure S11. Altered cell-cell communication signaling in P60 and P90 hNPC-treated RCS rat retina. (A)** Scatter plots showing Müller glia and hNPCs serving as one of the major sources and targets in hNPC-treated rat retina. **(C-J)** Network plots showing the strength of FGF **(C)**, PTN **(E)**, BMP **(G)** and JAM **(I)** signaling within different host retinal cell populations and with hNPCs in P60 and P90 hNPC-treated retina. Circle sizes proportionate to the number of cells in each cell group, and edge width represents the probability of communication. Violin plots showing the expression of ligands, and their receptors involved in FGF **(D)**, PTN **(F)**, BMP **(H)** and JAM **(J)** signaling in P60 and P90 hNPC-treated host retinal cells and grafted hNPCs.

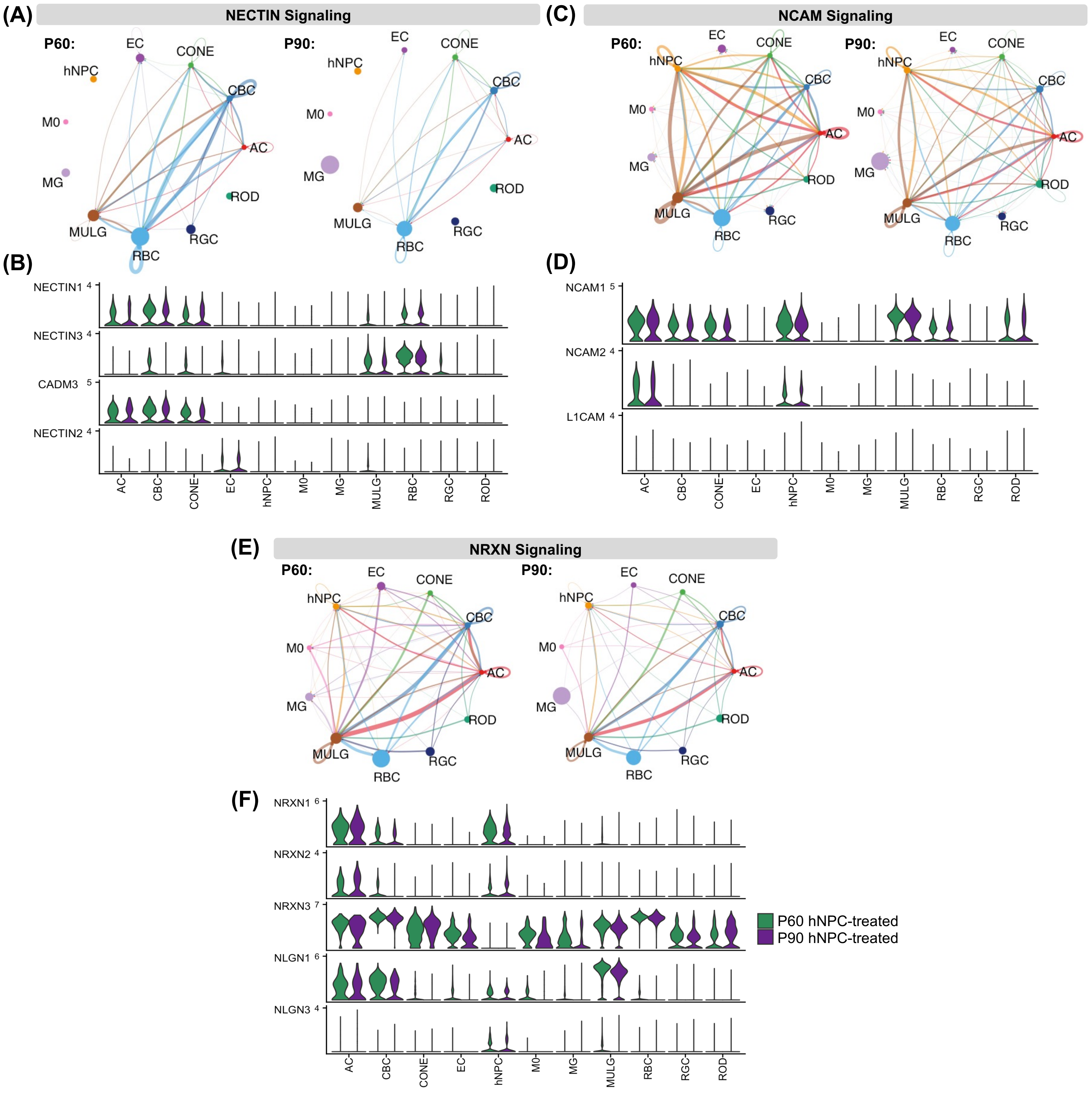


**Figure S12. Altered synaptic signaling-based cell-cell communication in P60 and P90 hNPC-treated RCS rat retina. (A-F)** Network plots showing the the strength of NECTIN **(A)**, NCAM **(C)**, and NRXN **(E)** signaling within different host retinal cell populations and with hNPCs in P60 and P90 hNPC-treated retina. Circle sizes proportionate to the number of cells in each cell group, and edge width represents the probability of communication. Violin plots showing the expression of ligands, and their receptors involved in NECTIN **(B),** NCAM **(D), and** NRXN **(F)** signaling in P60 and P90 hNPC-treated host retinal cells and grafted hNPCs.

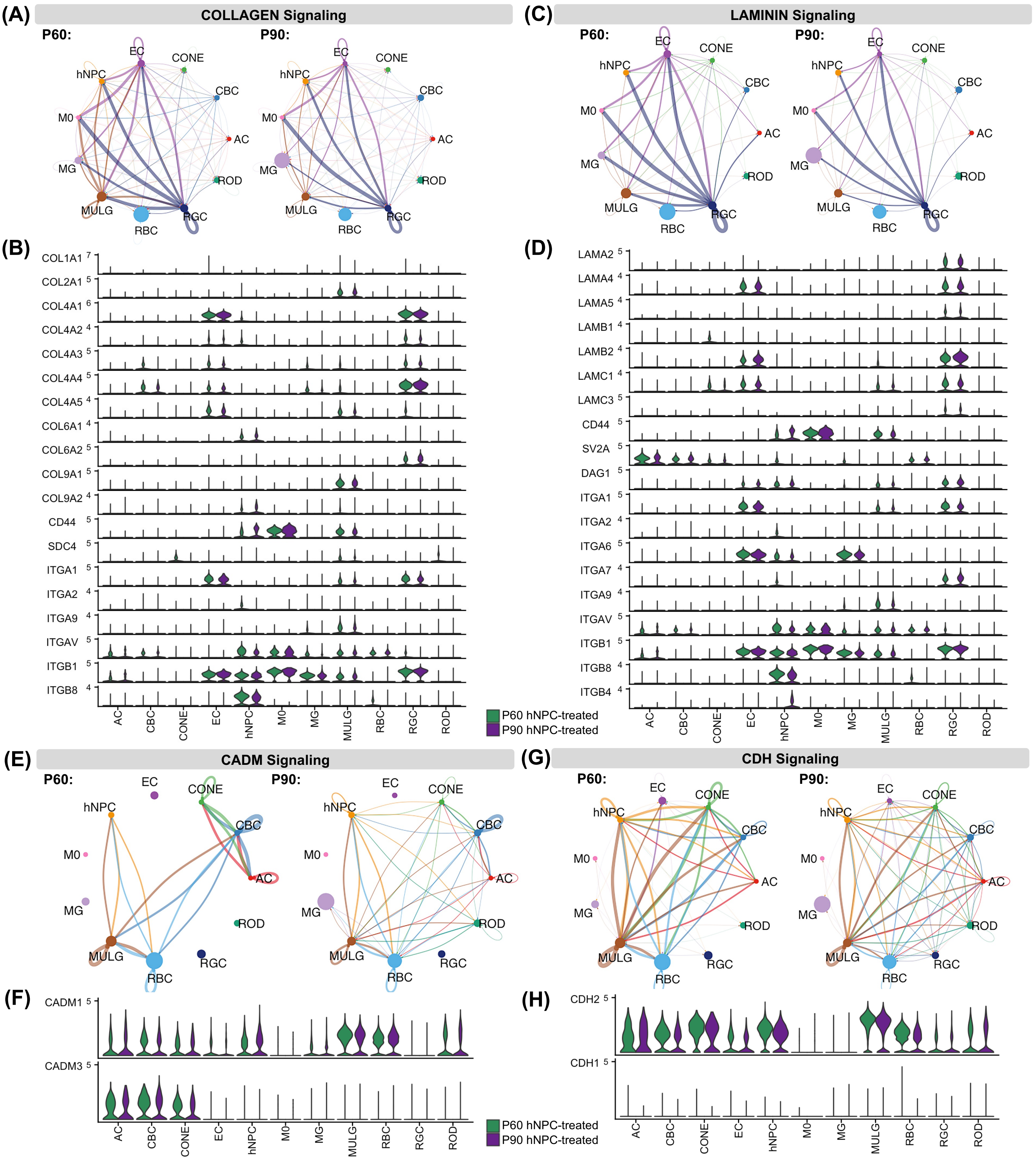


**Figure S13. Altered cell adhesion signaling-based cell-cell communication in P60 and P90 hNPC-treated RCS rat retina. (A-D)** Network plots showing the strength of COLLAGEN **(A)**, LAMININ **(C),** CADM **(E)**, and CDH **(G)** signaling within different host retinal cell populations and with hNPCs in P60 and P90 hNPC-treated retina. Circle sizes proportionate to the number of cells in each cell group, and edge width represents the probability of communication. Violin plots showing the expression of ligands, and their receptors involved in COLLAGEN **(B)**, LAMININ **(D),** CADM **(F)**, and CDH **(H)**, signaling in P60 and P90 hNPC-treated host retinal cells and grafted hNPCs.

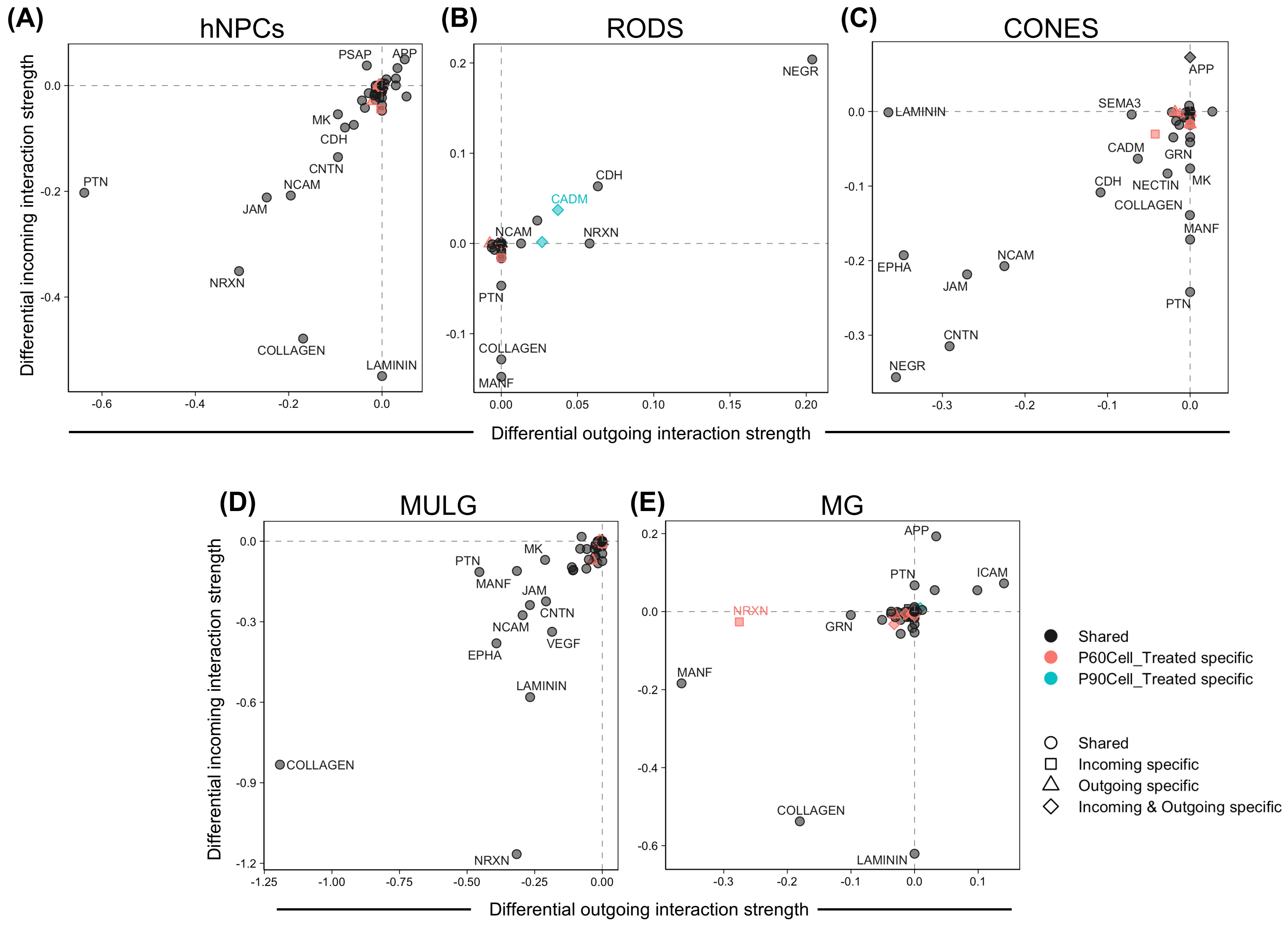


**Figure S14. Comparative differential cell-cell signaling analysis between P60 and P90 hNPC-treated RCS rat retina.** Scatter plots showing the differential outgoing interaction strengths versus differential incoming and interaction strengths from the differential cell-cell signaling network analysis between the P60 versus P90 hNPC-treated rat retina. In each cluster, the statistically significant ($p$-value $< 0.05$) signaling pathways were shown between P60 and P90 hNPC-treated RCS rat retina based on a permutation test.

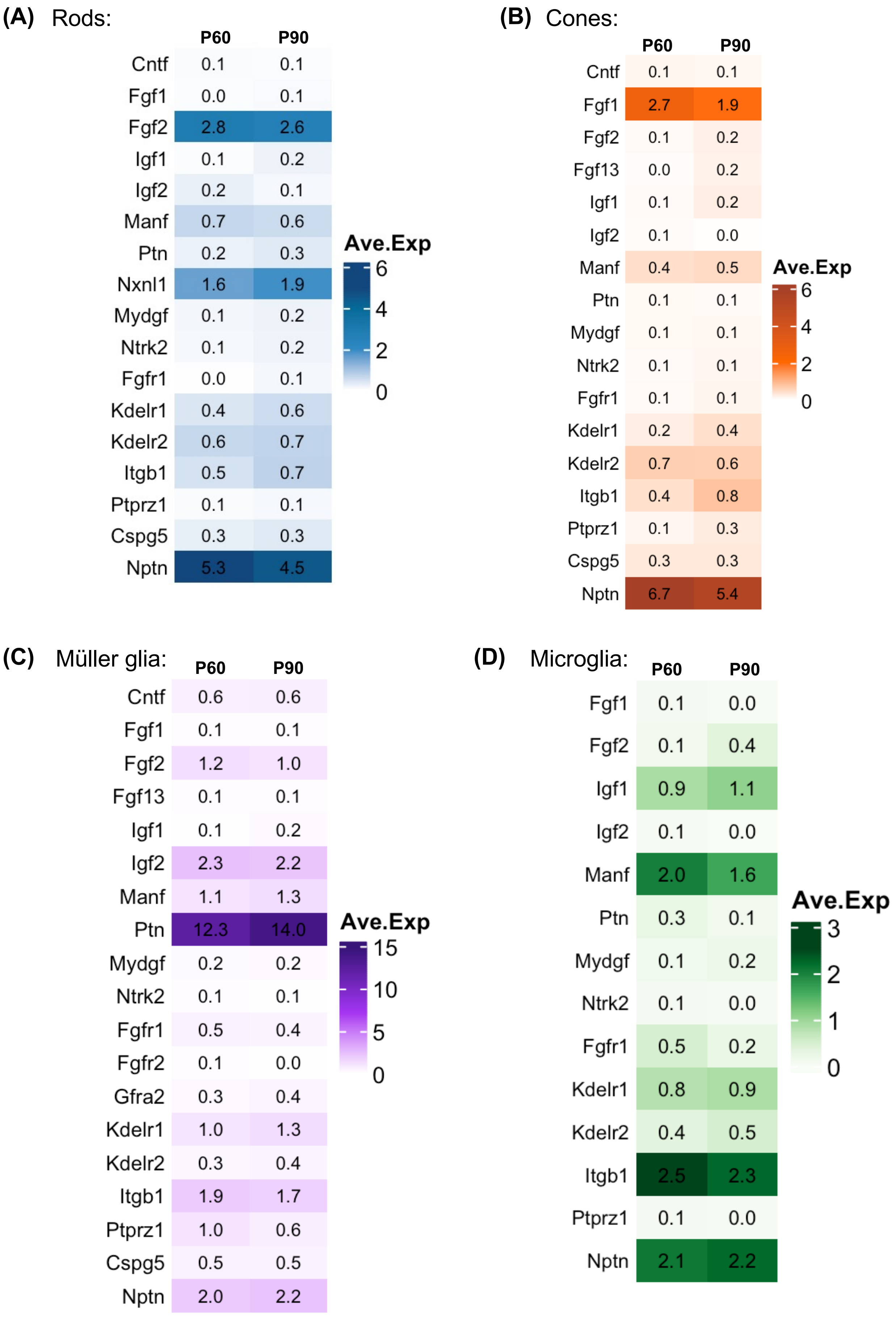


**Figure S15. Expression of neurotrophic and growth factors as well as their receptors in P60 and P90 hNPC-treated RCS rat retinal cells. (A-D)** Heatmaps showing the expression of different neurotrophic and growth factors as well as their receptors in P60 and P90 hNPC-treated rods **(A)** cones **(B)**, Müller glia **(C)**, and microglia **(D)**.
